# Molecular Biointerface Characterisation for an Implanted Medical Device Using Cryogenic Orbitrap Secondary Ion Mass Spectrometry (Cryo-OrbiSIMS)

**DOI:** 10.1101/2025.05.09.652996

**Authors:** Akmal H. Sabri, Kei F. Carver Wong, Anna Kotowska, Leanne Fisher, Jeni C Luckett, Jimiama Mafeni Mase, Lisa Kämmerling, Grazziela Figueredo, David J. Scurr, Amir M Ghaemmaghami, Morgan R. Alexander

## Abstract

Implanted medical devices often fail due to foreign body reaction (FBR), a process still not fully understood. This work presents a new depth profiling approach to provide insight into the spatial metabolomics of the biointerface of implants, revealing biomolecular strata representative of the host response. This study examines silicone rubber poly(dimethyl siloxane) catheters implanted in mice for 1 and 28 days. Cryo-OrbiSIMS was used in combination with ToF-SIMS to identify metabolite profiles from the biological deposit found on the implants after removal from the tissue which were previously unattainable using tissue sectioning. Machine learning and statistical analysis of the profiles were used to help identify early biointerface responses to the implant, including at 1 day of implantation the observation of elevated sugars and itaconate an immunomodulatory metabolite that modulates FBR. At day 28 inflammation associated markers were observed such as urate and palmitic acid (FA 16:0). Depth profiling revealed two distinct molecular layers in the deposits: amino acids, and nucleic acids were preferentially seen towards the host tissue, consistent with the observation of a cell monolayer in the tissue sections, whereas certain lipids and fatty acids where either at the catheter-deposit interface or towards the host tissue after 28 days. The stratification was less well developed at 1-day implantation, but common lipids were seen at the deposit-implant interface across both time points. These insights advance understanding of FBR and support the development of improved implant materials.

## Introduction

Medical devices play a pivotal role in modern medicine ranging from stents, catheters, hip and knee joints and sensors to treat and monitor medical conditions or replace and augment physiological functions [1]. Silicone-based biomaterials are commonly used for medical devices, applied as medical tubing, tissue replacement, intraocular lenses, voice prostheses, dialysis membranes, breast implants and ventriculoperitoneal shunts [1,2]. The response of the body to implants, also known as foreign body response, is described as complex multistage process characterised by an acute inflammatory phase which is the result of adsorption of biomolecules such as chemoattractant, cytokines, growth factors, and other bioactive agents leading to the formation of provisional matrix [3] [4]. This acute FBR phase is also characterised by the recruitment of immune cells such as neutrophils and monocytes to the site of implantation. As the response progresses, the protein within this provisional matrix undergoes a dynamic adsorption-desorption process, known as the Vroman effect resulting in the replacement of smaller proteins with larger ones. In addition, the recruited monocytes will undergo differentiation into macrophages at the site of implantation while releasing degrading enzymes and reactive oxygen species. During this period the fusion of macrophages into polynucleate foreign body giant cells occurs to phagocytose the implanted biomaterial. Overtime, this pro-inflammatory stage transitions into a chronic FBR response characterised by the presence of a fibrotic collagenous layer between the implant and the host tissue. As the FBR transition into the chronic phase the macrophages undergo a phenotypic switch from pro-inflammatory (M1) to an anti-inflammatory (M2). These M2 macrophages results in formation of a fibroblast and extracellular matrix-rich capsule that covers and isolates the implant. The surface properties of implant materials (topography, chemistry, compliance) play a critical role in modulating the foreign body response [5].

Schreib *et al* have proposed new mechanistic insights into the process of foreign body response to implanted medical devices [6]. Using Bi_3_^+^ Liquid metal ion gun (LMIG)/ToF-SIMS analysis, the researchers showed that lipid deposition on biomaterials after the first day of implantation, correlated with the implant’s immunogenicity which ultimately dictated the overall foreign body response to the implant. They were able to identify 11 fatty acids, using LMIG/ToF-SIMS analysis, that were enriched on the surface of explanted pro-fibrotic implants and uncoated PDMS discs, while phospholipid and sphingomyelin were enriched on the surface of anti-fibrotic implants. This correlation, and a potential role for specific lipids in controlling FBR, offers a novel adjunct to the orthodoxy of proteins being the main determinant of response to implants.

The OrbiSIMS has been developed with superior mass resolving power and accuracy, based on the Orbitrap analyser, along with softer fragmentation using argon gas cluster ion beams (GCIB) to achieve a more comprehensive interpretation of endogenous biomolecular processes, such as response of implants to host tissue, using SIMS [7,8]. Suvannapruk *et al.* applied GCIB/OrbiSIMS to the molecular characterisation of sections of tissue proximal to implants to provide information on the *in vivo* host response to medical device implantation, based on OrbiSIMS metabolite profiling of *in vitro* single macrophage phenotypes. The molecular characterisation of sections of tissue proximal to implants showed that monoacylglycerol (MG) and diacylglycerol (DG) secondary ions from the tissue were seen at increased intensity in the tissues ≈10 µm from the implants of reduced FBR collagen capsule thickness seen in histological sections.[8] Here, we take the approach of examining the surface of the removed implants using Cryo-OrbiSIMS, rather than analysing the histological sections, to provide insight into the biointerface.

The mass resolving power and accurate mass of OrbiSIMS datasets provide a rich but complex information source on the molecular and fragments from lipids, amino acids, and other small molecules. manual annotation of Orbi-SIMS data has been used to detect large range of biomolecular species ranging from lipids, amino acids, sugars, peptide fragments and proteins[9–11]. There is an impetus to utilise statistical algorithms, such as machine learning to aid in deconvoluting these complex datasets. The advancement new complex algorithms for artificial neural networks (ANN) and deep learning architectures underpinned advances in computational hardware [12]. For instance, several studies have recently been published that demonstrated the utility of machine learning in visualising and understanding hyperspectral SIMS datasets [13–15]. Recently, Bamford *et al* has successfully employed Self-Organizing Map and Relational Perspective Mapping unsupervised machine learning workflow on ToF-SIMS ion image to identify regions of extracellular vesicles at single pixel resolution released from microglia [15].

Here we apply a logistic regression analysis machine learning model coupled with a data-driven multivariate approach to identify and interpret biomolecules changes on the surface of the implanted silicone tubes. We probe the inflammatory phase by looking at tubes removed 1 day post implantation, and the early fibrotic phase using samples removed 28 days post implantation to elucidate the cascade of events that takes place during a foreign body response to a silicone implant. Automated peak assignment was achieved using molecular formula predictor (MFP) to identify the range of biomolecules deposited in combination with established databases[16]. It was found that cryogenic conditions augmented the signal of biologically relevant molecules from the surface of implants, consistent with previous reports [9,17,18]. This methodology and workflow are translatable to the analysis of novel biomaterials with immune-instructive properties in the search for materials capable of mitigating the FBR response *in vivo* and offering new insights into the molecular basis of the FBR.

## Materials and methods

### In vivo study

Male Balb/c mice (aged 56-62 days, approx. 25g) were used in this study. Mice were purchased from a recognised supplier (Charles River Laboratory, UK) and were initially be housed in groups of 5 (cage size 500 cm^2^ with sawdust bedding and sizzle nest) according to conditions stipulated by the Home Office. To acclimatise the animals to their surroundings, prior to experimentation, animals were housed for a period of at least 7 days without disturbance, other than to refresh their bedding and to replenish their food and water provisions. Each animal was randomly assigned a code which will dictate the implant (or implants) that it will receive and the form of processing to be undertaken at the time of harvesting. Animals will be anaesthetized using inhalation of isoflurane and air, and the dorsum of each mouse will be shaved and washed (0.05% aqueous Chlorhexidine Gluconate). The extremes of a 1.0cm line, dorso-ventral in orientation, will be “dot-marked” onto the shaved skin of each flank using a flexible transparent plastic template and an indelible marker pen. Seventy percent alcohol will be applied to the skin immediately prior to incision. A full-thickness incisional wound, 1.0cm in length, was created by incising the skin between these dot-marks. This incision will include the panniculus carnosus and hypodermis. The free edge of the skin will then be lifted, and one sub-cutaneous pocket will be created from each incision with the aid of blunt-nosed scissors. A single piece of catheter with an 12 mm external diameter and a length of 5 mm was wetted with Pharma grade sterile water and were inserted into each pocket. Care will be taken not to touch the face of implant materials. Following insertion, sites were inspected to ensure that samples are located appropriately. After sample insertion, the incisional wounds were closed using non-resorbable sutures.

After biomaterial implantation, animals were housed individually. Animals were maintained at an ambient temperature of 23°C with 12-hour light/dark cycles and were provided with food and water ad libitum. Following all anaesthetic events, animals were placed in a warm environment and were monitored until they have fully recovered from the procedure. All animals were provided with appropriate analgesia (Vetergesic, [buprenorphine]) after surgery and were provided additional analgesics as required. The mice were administered with prophylactic antibiotics (in the form of Enrofloxacin [Baytril], subcutaneous (s.c)) on the day of wounding and subsequently on every 7th post-operative day. All animal procedures were carried out in a Home Office licensed establishment under the following Home Office Licences (PCD: 50/2505; PPL: PP2870359; PIL: IBCEFDF55; PIL: I34817249). The health status of animals was monitored within two hours of implantation and subsequently daily for the first 7 days and weekly thereafter.

Animals were monitored daily for the first 7 days to ensure that sutures remain in place; sutures will be removed on day 7. Animals will then be maintained for a further 21 days until post-implantation day 28. At 1 day or 28 days post-implantation, animals will be humanely killed by a UK Home Office compliant method. Following confirmation of death, each animal will be rapidly dissected, and all implantation sites harvested. Excised implants will be snap-frozen in liquid nitrogen and shipped on dry-ice (-80°C). Implantation sites for histological investigation were fixed (10% neutral buffered formalin, Sigma, UK) and processed to paraffin wax. Excised tissue will be sandwiched between two pieces of foam, prior to being placed in fixative, to reduce the extent of tissue curling. Fixed specimens were trimmed and dissected in a cranio-caudal direction, generating two half sites (figure 3). The dorsal half were processed and embedded in paraffin wax (the remaining half will be stored in 70% EtOH). Specimens will be orientated in such a fashion as to ensure that appropriate transverse sections of the implantation sites were taken. 6µm sections were taken, processed and stained with H&E. In addition, the sections were stained using Picrosirius Red Stain (Polysciences, Hirschberg, Germany) and Masson’s Trichrome Stain (Polysciences, Hirschberg, Germany) kits. The image of the stained sections were acquired using Axioscan 7 from Zeis. This was done using tiled sequential acquisition with flash bright field at 20x objective. For section stained using Picrosirius Red, the image was also acquired using polarised light. Inflamed area measurement was done by taking macroscopic images of explanted skin tissue which were pinned down to flatten out the skin. Image J was utilised to analyse the area of inflammation surrounding the catheter segments by thresholding the colour change observed in the skin tissue.

### Sample preparation and Cryo-OrbiSIMS analysis

Excised implants were carefully removed by using a clear pair of forceps from the animal. A pair of isopropyl alcohol sterilised forceps were used to carefully used to grab the edge of the catheter during removal. This was done to avoid touching the middle of the cylindrical catheter as this will be the region by which analysis will be conducted. During this process the subcutaneous tissue that was attached to the skin was peeled from catheter at 180° axially to the tube, without touching the surface of the catheter. This revealed the strongly adhered biointerfacial deposit that have delaminated from the surrounding tissue. This surface was not washed or treated with any chemical and was immediately snap-frozen in liquid nitrogen and shipped on dry-ice (-80°C). The catheters were stored at -80^°^C until analysis. Prior to cryo-OrbiSIMS analysis, the sample were thawed at room temperature for 2 minutes. This is to enable the sample to be sliced in half and laid flat on a cryo-manipulation station. The implant consisted of PDMS catheter with a dimension of 12 mm external diameter and a length of 5 mm. The surface of catheter is smooth and consisted of a thin film of biological deposit with a watery appearance that was formed following implantation into the animal. To mount the catheter onto the cryo-stage, a pair of clean forceps were inserted into the internal diameter of the catheter to carefully hold the implant and a pair of sterilise scalpel were used to slice into the middle of the cylinder to cut the implant in halves. Both halves of the catheter was careful transfer onto the cryo stage by holding the edge of the catheter, this is done to done avoid contaminating the surface of the catheter prior to analysis. The samples were adhered to the cryo-manipulation station using a thin layer of OCT glue. Using a pair of forceps the samples were then snap freeze using liquid nitrogen. Before measurement by cryo-OrbiSIMS, samples were placed in a cryo-manipulation station, Leica EM VCM (Leica, Germany), from where they were transferred to the cryo-OrbiSIMS using a shuttle chamber Leica EM VCT500 (Leica, Germany).

The cryo-OrbiSIMS is equipped with a fully proportional–integral–derivative (PID) temperature controller, which controls resistive heating. This setup also incorporates a direct liquid nitrogen (LN2) closed loop circulation cooling stage to enable the sample to be thermoregulated under cryogenic condition within the load lock and main chamber. The instrument is installed with a cryogenic storage tank where LN2 was pumped for circulating the cooling medium through vacuum feedthroughs to a cooling finger below the sample, allowing fast cooling to −180 °C with a stability of ± 1–2 °C for at least 7 days. Mass calibration of the Q Exactive instrument was performed once a day using silver cluster ions. Electrons with an energy of 21 eV and a current of −10 μA and argon gas flooding were used for charge compensation. The surface of the catheter is curved, and analysis was conducted along the length of the catheter and analysis were avoided at the edge of the catheter where it was in contact with forceps during. ToF-SIMS analysis was conducted prior to OrbiSIMS data analysis. For ToF-SIMS images, the data were acquired using Bi_3_^+^ cluster source with a primary ion energy of 30 KeV was used, and the primary ion dose was preserved below 1 × 10^12^ per cm^2^ to ensure static conditions. The ToF-SIMS analysis was performed over an analysis area of 500 x 500 µm. For all Orbitrap data, mass spectral information was collected from a mass range from 80 to 1200 Da. The Orbitrap analyzer was operated in positive-ion and negative-ion mode at the 240,000 at m/z 200 mass-resolution setting (512 ms transient time). The GCIB OrbiSIMS resulting in consumption of the material on the surface of the catheter over 100 scans, which over a total ion dose of 1.29 x 10^14^. A total of n=30 technical replicates across N=3 animals per time point was acquired. The technical replicates per animal per time point was acquired along the length of catheter in order to capture as much chemical information that are present on the surface of the catheters. Each OrbiSIMS analysis was conducted using sawtooth raster mode with a crater size of 181.1×181.1 µm which equates to a field of analysis of 100 x 100 µm. Per time point, a total of n=30 technical replicates was acquired across N=3 animals, For replicate N1 for both time point, a total of n=6 technical replicate was run as this was the preliminary replicate. For N2 and N3 for both time point, a total of n=12 technical replicate was acquired.

### OrbiSIMS Peak Assignment

IonToF SurfaceLab 7.1.116182 were used to process the results and create the peak lists for export. Chemical filtering via SIMS-molecular formula predictor (MFP) was done *via* calculating the possible chemical formula permutation based on elemental restrictions from secondary ion data, including that obtained from a depth profile analysis. The elemental limitation applied for secondary ion assignment: C [4–150], H [2–250], N [0–30], O [0–20], S [0–1], P [0-2], Na [0–1], K [0–1] and Cl [0-1]. The double bond equivalence was set between -10 to 50. All possible chemical formulas were within the mass deviation of ± 5 ppm below m/z 95 and ± 2 ppm above m/z 95. All the predicted formula filtered based on the Human Metabolome and LIPID MAPS® databases.

Putative assignments were made using Human Metabolome Database (HMDB) filter based upon secondary ions accurate masses, *mz*. Lipid putative assignments were made using computationally generated bulk lipids database from LIPIDMAPS based upon secondary ions accurate masses, *mz*. These assignments are used to provide a subclass assignment, but the exact structures are not quoted since we do not have MS data. Some of these assignments may also result from fragmentation of larger species.

### Non-negative matrix factorisation

The IONTOF SurfaceLab® software was used to exports the data into a BIF 6 file format. For each dataset, Surface Lab 7.2 (IONTOF GmbH) was used to perform an automated peak search on the total spectra restricted only to peaks with intensity above 100 a.u. Peak intensities were then exported for each observation. Non-negative matrix factorisation (NMF) was performed using the simsMVA software. Prior to NMF, data were Poisson scaled to account for non-uniform noise across the mass spectra. NMF with 3 factors was achieved using a Poisson-based multiplicative update rule algorithm. The number of factors was chosen based on principal component analysis of a compressed matrix of the original dataset. A total of 500 iterations was conducted per dataset.

### Depth Profile Analysis Estimation

Information from published depth profiling of organic molecular standard (Irganox 1010) [19] was used to estimate the depth of GCIB depth profiling. This was based on information from primary ion dose of the GCIB, which is displayed as a primary X-axis in **Figure 6 (a)** and the depth is presented as a secondary x axis. The sputter yield from 20keV Ar_3000_^+^ on Irganox 1010 was adjusted for the cryogenic condition (-170°C) used in this study. This was done by comparing to the literature values using the calculations developed by Seah *et al*. [20] as used previously by Kotowska *et al* [21]. The depth profile was normalised to the maximum intensity of the each species in order for us to easily gauge the trend in profile of respective species as a function of depth. In addition, the profile was averaged over 10 scans per data point as previously reported[9,21], in order to improve the signal to noise ratio and gauge the trend in chemical species as a function of depth.

### Machine Learning of Cryo-OrbiSIMS data

Machine learning was explored to automate the discovery of deposited biomolecular species and to better understand the deposition within 1 day and 28 days of implantation. Two datasets of biomolecules deposition from the negative and positive ion spectra were analysed. Two datasets of biomolecules deposition from the negative and positive ion spectra were analysed, this consist of n=30 technical replicates across N=3 biological replicates per time point. Their biomolecular species intensities were identified using secondary ion mass spectrometry coupled to an Orbitrap mass analyser. The samples were scaled to intensity values between 0 and 1 to avoid numerical instability in the machine learning training process, where 0 represents the lowest intensity in the dataset and 1 represents the highest intensity. Machine learning approaches (Random Forest, SVM, XGBoost and logistic regression) were trained using Helix [22]. Logistic regression<u>—a more interpretable, linear machine learning (ML) and statistical approach—yielded</u> overall good fit, comparable to non-linear approaches and more interpretable results. It is a statistical method for predicting the probability of a binary outcome, such as the presence of 1 day and 28 days biomolecular species, based on predictor variables, such as the intensities of biomolecular species. It employs a multivariate linear regression function to learn the relationship between the outcome and predictors, followed by a logistic function (Sigmoid function) to map the learned relationship to a probability range of 0 to 1. This method is valuable for interpreting the impact of predictor variables, indicated by regression coefficients, especially in cases where the outcome is binary. To gain deeper insights into the variables identified as important by both linear and non-linear models, we also employed more complex, non-linear ML methods for comparison with logistic regression. Post hoc interpretability analyses <u>(using SHAP and another ensemble of interpretation approaches implemented in Helix</u>[23]) were then conducted to assess whether the salient features identified by logistic regression aligned with those revealed by non-linear models. Our intention was to either validate the initial findings or uncover supplementary information regarding OrbiSIMS-derived variables that exhibit non-linear associations with the outcome, thereby aiding in the discrimination between 24-hour and 28-day profiles. To facilitate the identification of the most informative predictors, we applied Least Absolute Shrinkage and Selection Operator (LASSO) regularisation prior to model development for variable selection. This procedure reduced the dimensionality of the input space by limiting the number of independent variables, thus mitigating the risk of overfitting across both linear and non-linear models. Evaluation on an independent test set—following cross-validation on the training set—indicates that the models generalise well, suggesting that overfitting is unlikely to have occurred. Additional details on the machine learning methodologies and output are detailed in the supplementary information.

## Results and Discussion

### Histology

Sections of silicone catheters were implanted subcutaneously into male Balb/c mice as shown in **Figure 1 (a)**. Upon implant removal after 1 day, there were no visible differences in the surrounding tissue as seen in **Figure 1 (b)** but a transparent film was apparent on the surface of the material. After 28 days of implantation, a significant area of reddening was observed in the surrounding tissue, as well as the presence of vascularization believed to be indicative of prolonged inflammation at the site. This reddened tissue area was quantified in **Figure 1 (c)** by using Image J to analyse the area of inflammation surrounding the catheter segments by thresholding the colour change observed in the skin tissue. Histology performed on the tissue both surrounding and within the catheter tube are shown in **Figure 1 (d)**. The catheter have been removed post fixation and prior to embedding during sample processing. The tissue surrounding the implants after 28 days were stained using 3 separate complementary approaches: haematoxylin and eosin (H&E), Masson’s trichrome (MT) and Picrosirius red stains (bright field and polarised light) in order to get a full characterisation of the tissue. The results of all three stain are shown in **Figure S1**. A fibrotic layer is clearly indicated by the MT stain in **Figure 1 (d)** around the catheter at 28 days consistent with a FBR [24]. The polarised Picrosirius red stain along with the Masson’s trichrome stain clearly reveals a cellular layer on the implant side of the fibrous layer **Figure 1 (e)**. Previous report[25] have considered these two layers as one collective strata, i.e. the entire fibrotic biolayer, however these images show that this can be further subcategorized. The cellular layer was measure using 7 samples as 6 µm ± 1 µm and the collagen layer was which was 16 µm ± 5 µm **Figure 1 (e)**.

**Figure 1.**
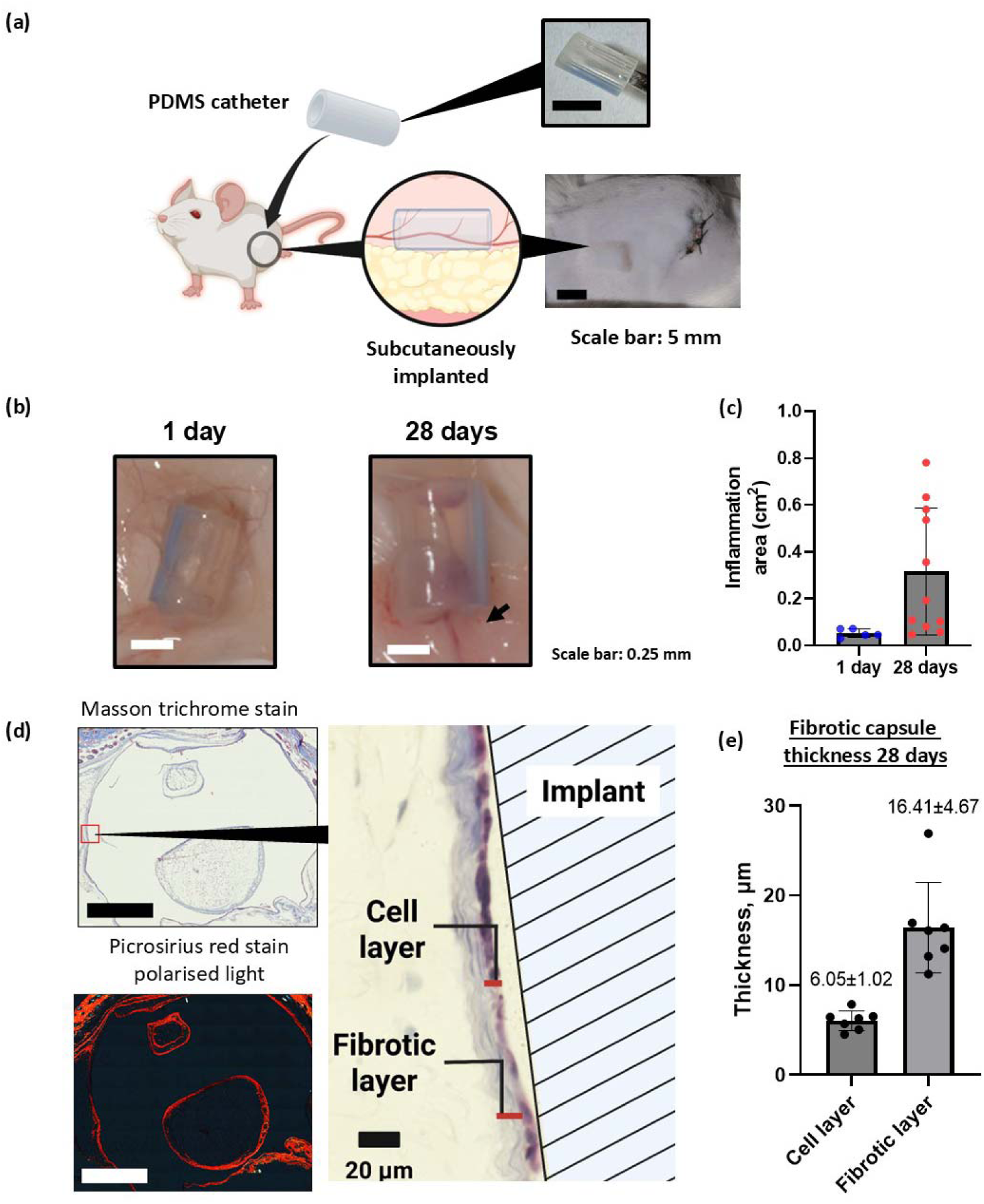
**(a)** Schematic illustrating the subcutaneous implantation of silicone catheter into male Balb/c mice. **(b)** Images of the implanted catheter in the subcutaneous layer of mice at 1 day and 28 days, scale bar: 0.25 mm. **(c)** Area of reddened tissue (indicated by arrow) showing inflammation surrounding the catheter following implantation along with the presence of vascularisation. Data are expressed as mean ± SD (n=5 for 1 day, n=11 for 28 days). **(d)** Masson trichrome and polarised Picrosirius Red staining and of FBR tissue sections at 28 days, scale bar: 1 mm. **(e)** Cellular and collagen layer thickness measured from histological sections.

### Biointerfacial chemical analysis

To study the molecular response of the host to the implant at an interfacial level, GCIB OrbiSIMS analysis of the samples was conducted on the surface of the explanted silicone catheter resulting in consumption of the material at the interface over a 100 analysis scans. Samples need to be examined either frozen hydrated using cryogenics or dehydrated at room temperature to allow them to be placed under vacuum for SIMS analysis. The implant consisted of PDMS catheter with a dimension of 12 mm external diameter and a length of 5 mm. The surface of the removed catheter was visually smooth with a watery appearance upon removal from the tissue. To mount the catheter onto the cryo-stage, a pair of clean forceps were inserted into the internal diameter of the catheter to carefully hold the implant and a pair of sterilise scalpel were used to slice into the middle of the cylinder to cut the implant in halves. The half catheter was careful transfer onto the cryo stage but holding the edge of the catheter. The surface of the catheter is curved, and analysis was conducted along the length of the catheter and analysis were avoided at the edge of the catheter where it was in contact with forceps during transfer.

We compared these two approaches; the physical effect of the sample drying under vacuum at room temperature was seen as cracks in the LMIG/ToF-SIMS images in **Figure 2(a),** where the water evaporation has led to shrinkage and fracture of the biomolecular layer. When the samples were analysed under cryogenic frozen-hydrated conditions, no cracking was observed. Plotting the intensities of the ions acquired from the room temperature samples against those acquired under cryogenic conditions in **Figure 2 (b),** clearly indicates that employing cryo-GCIB/OrbiSIMS augments the ion signal intensity for approximately 90% of ions, some by as much as 10^4^. Furthermore, more than 40 biomolecular species were only detected under cryo, including ions putatively assigned to; a diacyl-glycerophosphoethanolamine, PE 36:1, a diacyl-glycerophosphoinositol, PI 38:5, Glutamine and the oxidised fatty acyl molecules such as NAT 32:0;O4 and NAT 32:7;O. The augmented ionization yield is attributed to the presence of water that serves as a matrix to provide H^+^ transfer for secondary ion formation. Furthermore, cryogenic analysis prevents loss of molecules that are prone to evaporation such as fatty acyls, so frozen hydrated acquisition is adopted for the rest of the work in this paper [9,17]. It has recently been shown that performing OrbiSIMS analysis under cryogenic conditions enhances molecular stability during SIMS analysis by reducing molecular fragmentation and suppressed ion beam-induced damage, in agreement with previous literature [17].Cryogenic conditions can introduce artifacts such as ice crystal formation, altered ionization efficiency, or redistribution of analytes [26]. This is especially true for thick biological tissue or cross-sections, where the presence of ice crystals due to slow freezing can introduce fissures and holes in the specimen due to expansion of water upon freezing. In our study, we took several measures to mitigate these issues. The samples consisted of a very thin layer of biological deposits that was rapidly frozen using plunge freezing to minimize ice crystal formation and preserve native spatial distribution.

**Figure 2.**
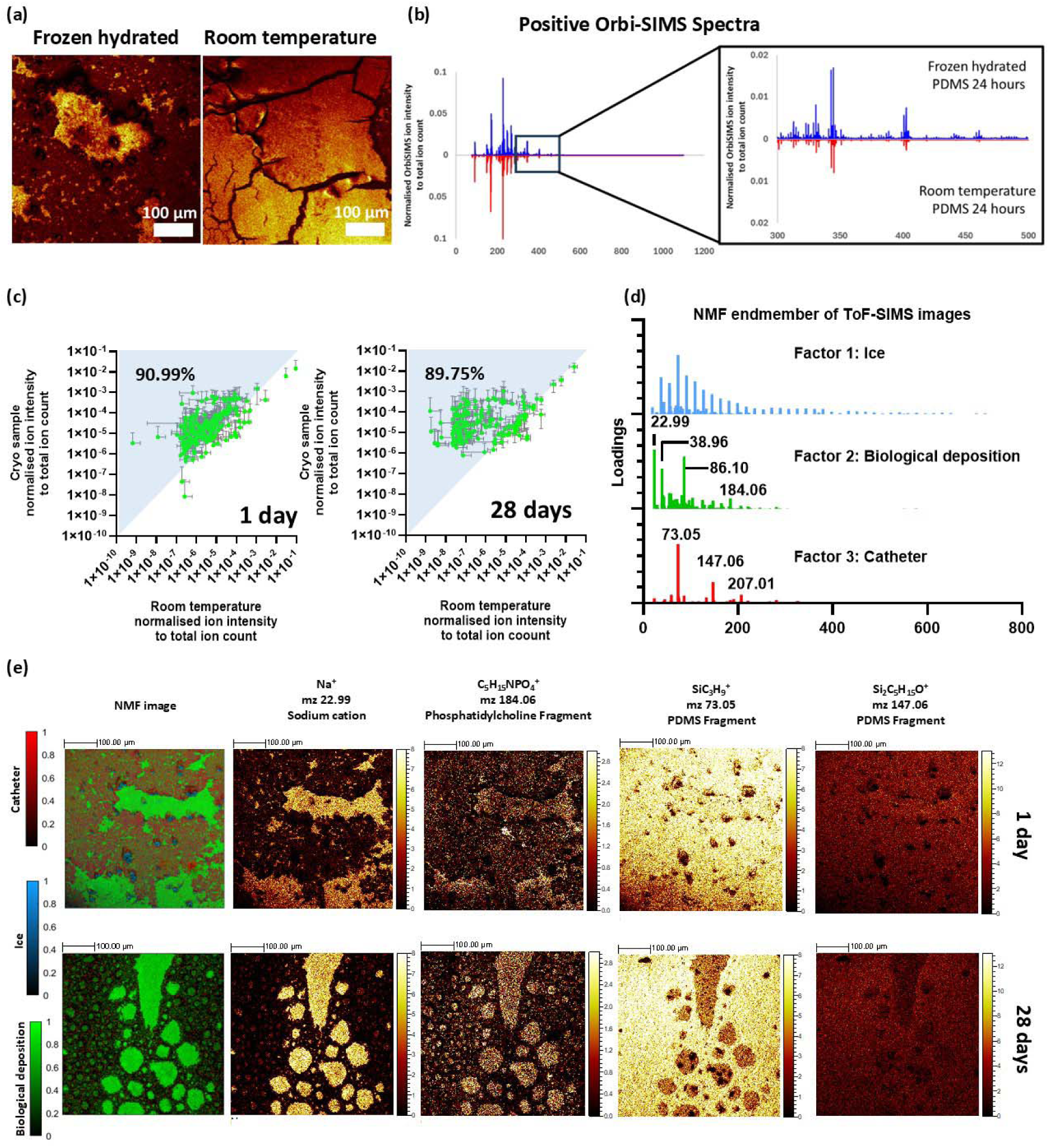
**(a)** LMIG/ToF-SIMS ion images of the catheter surfaces when sample were analysed under frozen hydrated and room temperature conditions. **(b)** Comparison of GCIB/OrbiSIMS spectra of catheter surfaces analysed under frozen-hydrated (blue) and room temperature (red) conditions. **(c)** Scatter plot comparing the normalised ion intensities of biomolecules detected when the surface of the catheters was analysed under room temperature and frozen hydrated. **(d)** Non-negative matrix factor (NMF) loadings showing the separation of LMIG/ToF-SIMS data into three distinct components: ice, biological deposition, and catheter with **(e)** representative NMF images for 1- and 28-days implantation following 500 NMF iterations. The NMF analysis was performed using loadings and images from raw LMIG/ToF datasets. Colour scale represents the intensity of NMF endmembers post calculation along with the ion images for C_5_H_15_NPO ^+^ which represent phosphatidylcholine fragments and SiC_3_H ^+^ and Si C H O^+^ which is used as a marker for PDMS. Scale bar: 100 µm. A red-green colour-blind alternative of Figure 2(e) is also available in **Figure S9** in supplementary information.

To enable the interpretation of imaging mass spectral datasets generated in ToF-SIMS analysis, non-negative matrix factorisation (NMF)[27] analysis was employed as shown in **Figure 2 (d)**. This revealed that the surface of the segments were very chemically heterogenous, as shown in **Figure 2 (e)**, with patchy biological deposit on the silicone tube. NMF images categorised the lateral chemical variance into three distinct surface chemistries along the surface of the catheter: secondary ion fragment representing the silicone from the catheter, ice and biological deposit at 1 day and 28 days. Contributions to these were ions representative of the silicone catheter and mobile oligomers using *m/z* 73.05 (SiC_3_H_9_^+^), 147.06 (Si_2_C_5_H_15_O^+^) and 207.03 (Si_3_C_5_H_15_O_3_^+^), ice where water clusters H_7_O_3_^+^, H_9_O_4_^+^ H_11_O_5_^+^ from H_2n+1_O_n_^+^ [28] were observed, and biological deposits containing phosphatidylcholine fragment ions and salts; 86.10 (C_5_H_12_N^+^) and 184.06 (C_5_H_15_NPO_4_^+^) 22.99 (Na^+^), 38.96 (K^+^). In **Figure 2 (e) and Figure S11,** it is apartment that there was a greater coverage of biological deposit on the silicone surface at 28 days relative to 1 day. Displaying selected individual ion images in **Figure 2 (e)** shows that biological deposit sits as discontinuous patches on the catheter (m/z 73.05) at 1 day. At 28 days large regions of biological deposit are seen as islands, which on some replicates becomes the dominant continuous phase in Figure S11/S12 replicate 1. The biodeposit phase represented by C_5_H_15_NPO_4_^+^ at the surface of the PDMS arises from the delamination of the tissues during the 180°peel of the tissue away from the catheter which exposed this interface critical to gaining insight into the response of the host to the implant.

### Cryo-OrbiSIMS biomolecular deposit spectral annotation

The Human Metabolome and computational LIPID MAPS® databases was used to help assigning the peaks from the OrbiSIMS analysis. Including [M+Na]^+^, [M+K]^+^, and [M+Cl]^-^were used to undertake an untargeted analysis of the biomolecular deposit captured in the cryo-OrbiSIMS spectra using the molecular formula prediction (MFP) approach. [16] The peak search of the GCIB/OrbiSIMS spectra resulting in identification of 4922 secondary ion peaks in the negative polarity and 1458 peaks in the positive polarity **Figure 3(b).** From this, we were able to putatively assign 90 lipid molecular ion peaks, 17 of which are fatty acids peaks. It is likely that these fatty acid peaks may arise as fragment ions from larger lipid molecular ion peak. In addition, were able to assign 214 metabolites molecular ions on the surface of the silicone catheters using the HMDB filter. Consistent with the greater area of biological deposit observed in the ToF-SIMS images, it was observed that there were more metabolites and lipids were detected on the surface of the tube at 28 days relative to 1 day.

**Figure 3.**
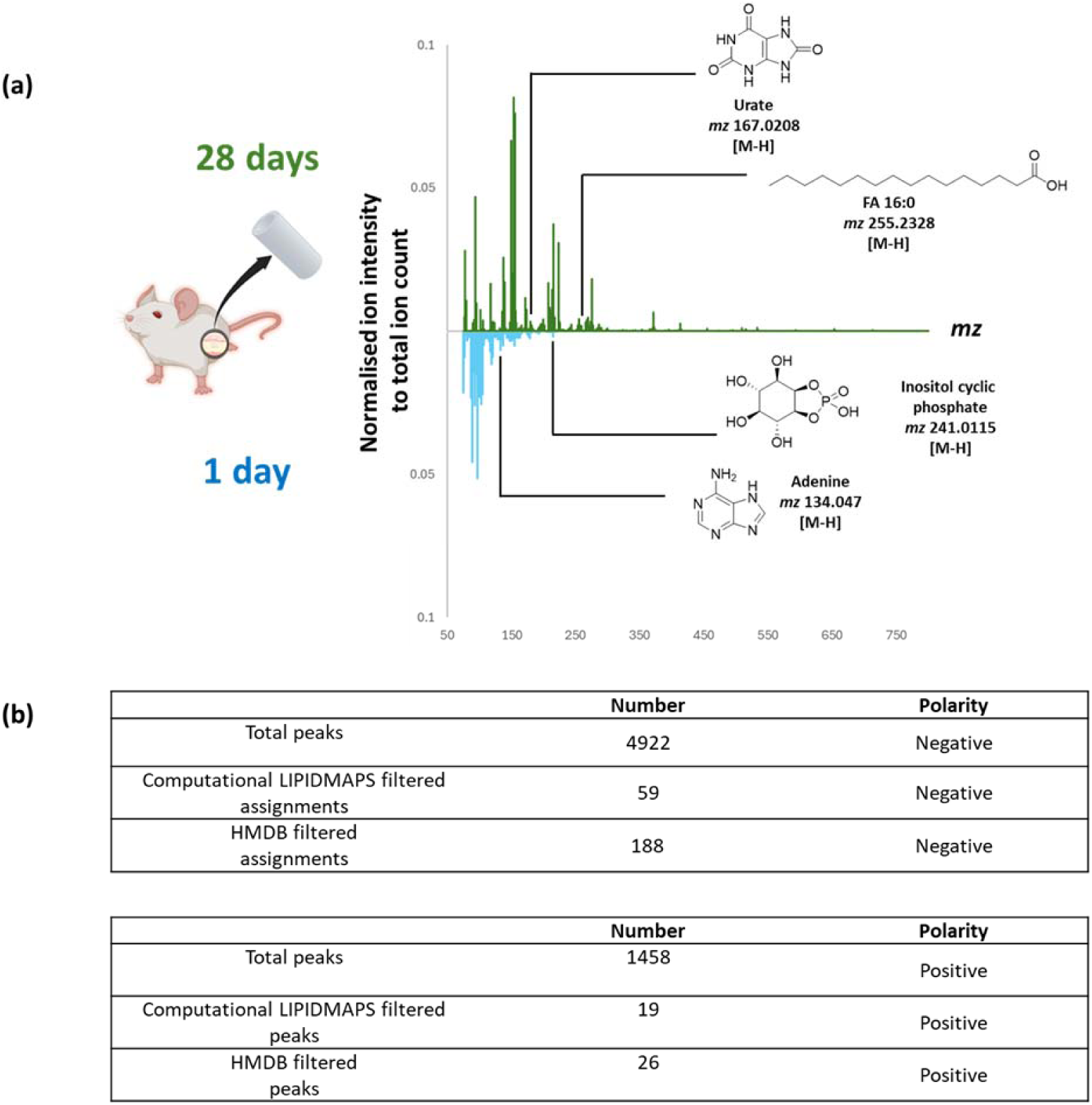
**(a)** Comparison of cryo GCIB/Orbi-SIMS spectra of silicone catheter surface at 1 day and 28 days **(b)** Table summarising the total number of peaks detected across the catheter surface at 1 day and 28 days along with the number of lipid and metabolite unique peaks.

### Identifying significant assignments using a Logistic Regression Machine Learning algorithm

To determine if we could better extract data from the OrbiSIMS datasets of the whole biointerface with machine learning, we employed a linear logistic regression and other non-linear machine learning models, including Random Forest, Support Vector Machine and Extreme Gradient Boost (XGBoost) on the assigned metabolites illustrated in **Figure 4 (a)**. The model aims to predict the difference between two classes (samples measured after 1 day or 28 days post implantation). The machine learning model was applied to all putatively assigned biomolecular species in each condition and from both positive and negative polarity combined. The details of the methodology and output of all the machine learning models that were ran for these datasets are detailed in the Supplementary Information (**Figure S2-S6** and **Table S3-S5**). Overall, the results throughout all models were consistent, with the best results obtained by Logistic Regression, using 5-fold cross validation and 20% of the data as a holdout test set. The workflow highlighted in **Figure 4 (a)** showed that the supervised machine learning method using logistic regression. The logistic regression-based machine learning model highlighted that there were 54 assigned species that were critical in differentiating the interfacial response between explanted catheters at 1 day and 28 days. The statistical significance of these metabolites was tested using a simple unpaired Student’s T-test and tabulated in **Figure 4(b).** This analysis revealed that 79.6 % of the species identified by the machine learning model were statistically significant, *p-value* < 0.05. The tabulated metabolites in **Figure 4(b)** also displayed mean coefficients values, with negative values being more associated with metabolites that are predictive of 1 day response while the more positive coefficient are associated with 28-day response. Interestingly, the top two phenotypic metabolite predictors for 1-day post-implantation are the glycerophosphoglycerol PG 39:1;O and linoleic acid FA 18:2 that suggest that these lipids are strongly associated with the surface of the catheter upon initial implantation or may be continually produced by the cells associated with the implant surface. These lipids may form the initial lipid layer on the surface of the catheter, for example from local cell extracellular vesicles as proposed by Screib et al.[29].

**Figure 4.**
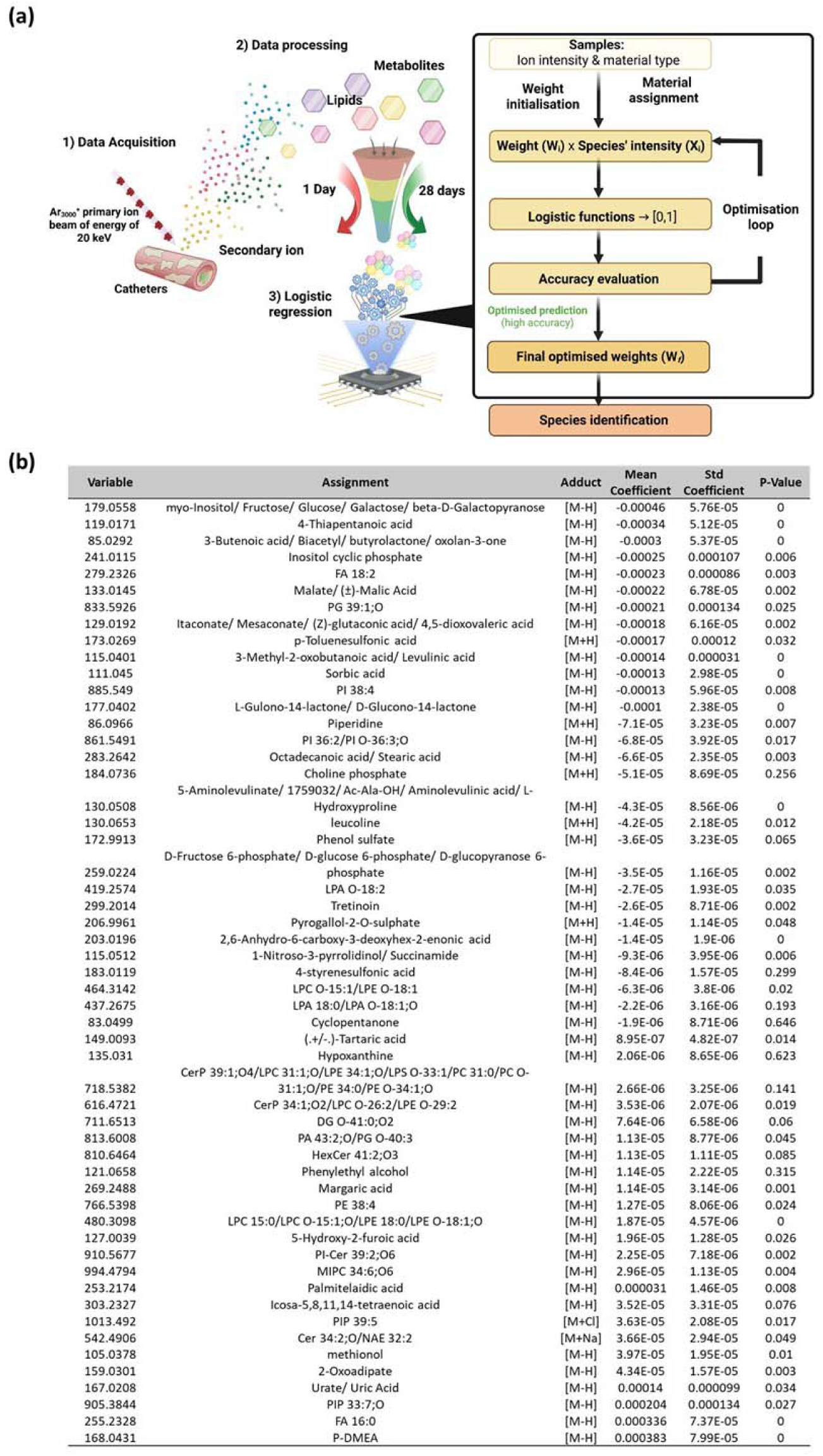
(a) Schematic of the data analysis workflow along logistic regression-based machine learning to identify metabolites that are important in differentiating implanted catheters at 1 day and 28 days. (b) Table indicating logistic regression average coefficients and statistical significance of Human Metabolome Database filtered dataset and computational LipidMAPS filtered dataset that are important in differentiating the surface chemistry of the catheters at 1 day and 28 days. Negative mean coefficient indicates metabolites that are more associated with catheter surface at 1 day while positive mean coefficient indicates metabolites that are more associated with catheter surface at 28 days.

### Fold change and statistical significance

In addition to machine learning, we have used a volcano plot to help interpret the complex Orbi-SIMS dataset from the biointerface. These are useful in comparing two sample types with a large number of data components, replicates and dynamic range. We display the mean ion intensity from n=30 replicate analyses on a plot of log_10_ *p-*value determined between the two implant time points versus the fold change over time on a log_2_ scale in **Figure 5 (a).** Secondary ions were normalised to the MFP derived species for their respective group total. This was done to minimise the impact of any variability in secondary ions between samples from other chemical species [30,31]. It was observed that there were 61 metabolites that are of statistically significant higher concentration (*p* <0.05, fold change ≥ 2) at 1 day. These include myo-inositol and several sugar isomers. For 28 days implantation we observed 15 metabolites that are expressed at a statistically significant higher concentration that include known markers of inflammation such as FA 14:0 [32]. It was observed that the urate levels on the surface of the catheters from 1 day to 28 days was seen to remain the same (*p*>0.05), suggesting sustained inflammation in the tissues surrounding the catheter. Urate is known end product for the metabolism of purines, the main constituents of nucleotides [33]. During the FBR cascade, urate particularly in it’s crystalline form has been reported to promote a pro-inflammatory response through the innate immune system, leading to inflammation and potentially tissue damage [34].

**Figure 5.**
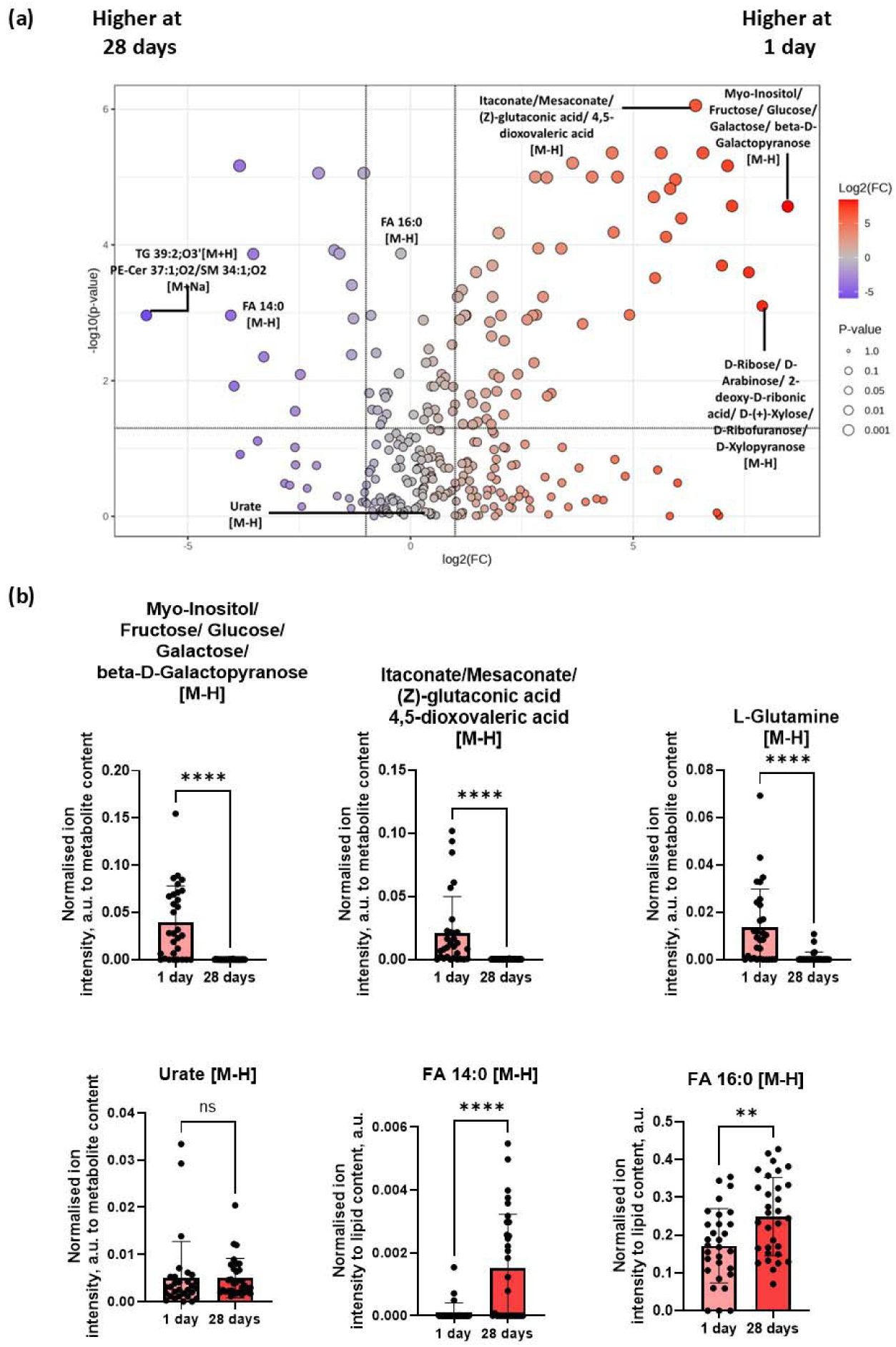
(a) Volcano plot of biomolecular species detected on the surface of catheters at 1 day and 28 days. Highlighted in the plot are some of the species that displayed a two-fold difference in species concentration with a statistical significance of *p* < 0.05. The volcano plot was constructed using FDR-corrected p-values from a t-test to determine statistical significance. For species plotted, the data are expressed as the average of all technical replicates, n=30 (b) Bar charts of selected biomolecules. Data are plotted as normalised ion intensities to total metabolite/lipid content. Data are displayed as n=30, mean ± SD, following non-parametric T-test with a statistical significance of *p* < 0.05. Non-parametric T-test was chosen due to the data did not display a normal distribution following Shapiro-Wilk and Kolgomorov-Smirnov tests. The output of the normality tests is detailed in **Table S3**.

Sugars such as fructose and glucose as well as metabolites that play key regulatory role in glycolysis such as itaconate and mesaconate [35,36] are found to be significantly higher on the surface of the silicone catheter on day 1 relative to 28 days in **Figure 5 (a)-(b)**. These biomolecules were also identified by the logistic regression-based machine learning algorithm to be predictive of phenotypic response towards silicone catheter surfaces at 1 day as shown in **Figure 4 (b)**. This observation is consistent with the kinetics of an inflammatory response [37]. During the early phase on an inflammatory response the damage tissue and its periphery, in this case the surface of the catheter, will experience a burst of itaconate. This molecule has been extensively reported to play a role in regulating and limiting the local inflammatory response which in turn prevent further damage to the tissue should there be any sustained inflammation [38,39]. Itaconate is produced by activated macrophage and serves as an immunometabolite that modulates FBR by inhibiting succinate dehydrogenase while simultaneously activating NRF2 and ATF3 [40]. These effects collectively suppress NF-kB signalling which reduce pro-fibrotic macrophage activation. The immunomodulatory role of itaconate was further reinforced by a recent work that showed implants that are loaded with itaconate and citrate displayed reduced foreign body response on the surface of implanted cardiac patches [41]. In addition, mesaconate and itaconate are constitutional (structural) isomers, since they share same molecular formular but differ in atomic connectivity. Both molecules are related dicarboxylic acids and are both intermediates in metabolic pathways, particularly in microbial and mammalian system. Despite these differences they displayed similar anti-inflammatory effects, possibly *via* overlapping the NRF2 activation and cytokine suppression pathway [35].As the inflammation subsides, itaconate levels also decrease, as shown in our analysis at 28 days in **Figure 5(b),** reflecting the reduced need for immune suppression. Overall, both itaconate and mesaconate are emerging immunomodulatory metabolite, with growing recognition of their roles in the cascade of immune regulatory events during the foreign body response.

The sugar molecules identified in the analysis are linked to inflammatory processes particularly in macrophages through the pentose phosphate pathway or *via* glycolysis [42]. Viola *et al* and Liu *et al* have discussed how macrophages switch to glycolysis and pentose phosphate pathway to produce energy which helps augment cellular pro-inflammatory responses during an inflammatory response [36,43]. In addition, it has also been shown recently how both itaconate and mesaconate exert anti-inflammatory effects in pro-inflammatory macrophages as well as playing a role in inhibiting glycolysis [35]. A significant decrease in L-glutamine on the surface of the catheter from 1 day to 28 days was seen in **Figure 5 (b),** suggesting the utilisation of the amino acids *via* catabolic processes that drives inflammation. L-glutamine which is one of the most abundant amino acids in the human body, has also been linked to inflammation *via* glutaminolysis [44]. An increase in saturated fatty acids deposition, FA 14:0 and FA 16:0, was highlighted at 28 days relative to 1 day. These fatty acids have been studied extensively and have been shown to be linked to an increase in inflammation [45,46] as well as lipid-induced cellular apoptosis [47,48]. FA 16:0, also known as palmitic acid, is not inherently a TLR agonist. However, upon entering cells FA 16:0 can be converted into phospholipids, diacylglycerol and ceramides which then leads to activation of various signalling pathways that are common for LPS-mediated TLR4 activation, culminating in an inflammatory response. In particular, metabolic products of palmitic acid affect the activation of various protein kinases C, endoplasmic reticulum stress and cause an increase in ROS generation that promotes further inflammation [49]. In addition, FA 14:0 which is also known as myristic acid have been reported to exacerbate and promote further inflammation particularly in the presence of FA 16:0 [50,51].

### Comparison of data mining methods

A comparison of utility of the machine learning and volcano plot approaches in identifying the species associated with the catheter at 1 day and 28 days, shows that up to 37% of the metabolites identified by logistic regression-based machine learning overlapped with the conventional univariate approach in **Figure S2**. The ensemble of both methodologies, by machine learning and univariate statistical analysis increases the confidence in the results, as multiple perspectives of importance of the variables are taken into consideration, both looking at individual importance but also how variables can be important synergistically. Each methodology identified unique species, with the machine learning highlighting a narrower selection of metabolites relative to volcano plot analysis. The narrower metabolite list is attributed to Least Absolute Shrinkage and Selection Operator (LASSO) feature selection prior to machine learning. LASSO zeros the beta coefficients of those variables that are not significant to the outcome after which logistic regression-based machine learning was performed on the with the 54 variables to enable the metabolite to be ranked based on their contribution at 1 day and 28 day. This feature removal obviates the contribution of metabolites that had very little influence in the ranking. Since there are potentially useful species identified uniquely by each approach, we propose that they should be used in concert to help deconvolute the complex OrbiSIMS dataset.

### Molecular depth profiling and biointerfacial characterisation using OrbiSIMS

To elucidate the identity of this biointerfacial deposit as a function of depth, a GCIB/OrbiSIMS depth profiles were conducted shown for selected ions in **Figure 6 (a)**. Metabolites that showed the highest signal intensity at the start of the profile and decreases as a function of dose of primary ion beam and depth are categorised as species which were next to the cells in the H&E sections before removal from the tissue (**Figure 1d**), these are known as metabolite located at cellular interface. Species that display higher intensity later in the depth profile are categorised to originate from the material-host interface at the implant surface, these are known as metabolite distant from cellular interface. Non-lipid metabolites such amino acids and nucleic acid bases were concentrated at the at the surface of the deposit which are in close contact with the host tissue interface. In contrast, lipids such diacylglycerophosphoglycerol, PG 39:1;O and oxidised glycosphingolipids, SHexCer 41:5;O5 are buried deeper in the deposit and are located more closely with the surface of the implant itself. The putative structures of these metabolites identified in the depth profile are shown in **Figure 6 (c)**.

**Figure 6.**
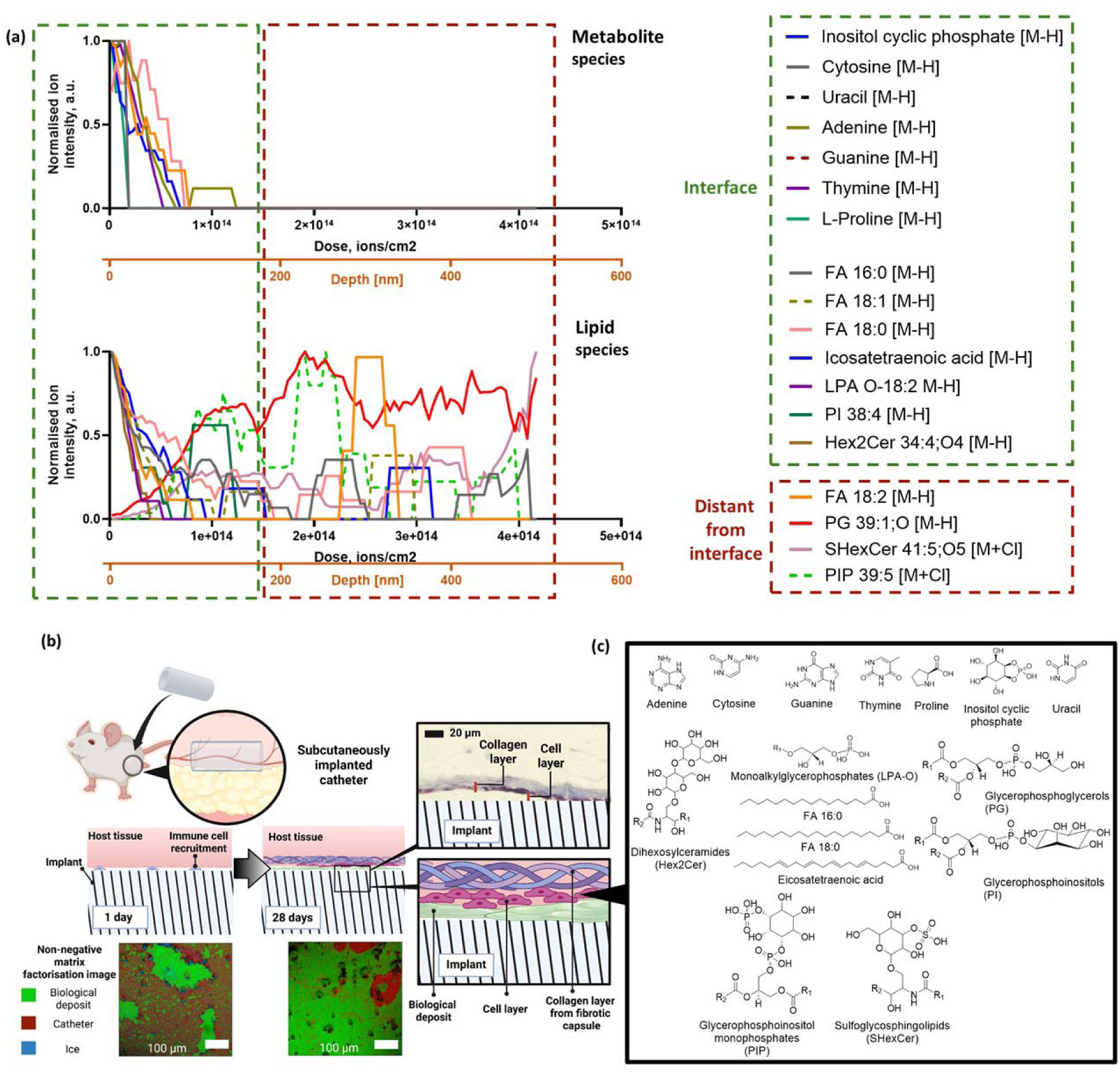
**(a)** Depth profile of silicone catheter surfaces implanted for 28 days versus primary ions dose (ions/cm^2^) and estimated depth displaying lipids, amino acid and nucleic acid bases molecular ions. Molecules present at the interface of the deposit, and the tissue (close to the surface) are categorised as interfacial while those present deep in the deposit away from the surface are categorised as distant from interface. The secondary x-axis presents depth estimated by comparison with organic standards **(b)** Schematic illustrating and summarising the presence of biological deposit formed on the surface of medical device following implantation **(c)** Chemical structure of species detected by cryo-OrbiSIMS. A red-green colour-blind alternative of Figure 2(e) is also available in **Figure S10** in supplementary information.

The surface of the explanted tubes, which was next to the cellular layer seen in the histology, was characterised by a number of metabolites, thymine, uracil, adenine, cytosine, guanine, L-proline and inositol cyclic phosphate, which we assume are related to cells. Some lipids were also found at this interface, include icosatetraenoic acid and the glycerophosphoinositol PI (38:4). These lipids classes have been shown to be involved in a variety of biological processes as well as being a key marker for plasma membrane and endoplasmic reticulum [52,53]. It was seen in **Figure 4 (b)** that oxidised diacylglycerophosphoglycerol, PG 39:1;O and fatty acid, linoleic acid FA 18:2 species were ranked by logistic regression-based machine learning algorithm as biomolecules that are mostly associated with the surface of the implants at 1 day. It can therefore be postulated that these lipid species may form an initial lipid deposit that is closely associated with the surface of the catheter upon implantation. The presence of these initial lipids on the surface of the catheter may arise from the deposition of lipid from surrounding tissue due to initial implantation procedure. Overtime other lipid species and metabolites are deposited onto the lipid layers resulting in the formation of a stratified lipid deposit layers. Overall, this work showed that the use of cryo-OrbiSIMS and machine learning provides us the tools needed to uncover the different layers of lipids and metabolites that make up the biointerface which reflects the molecular history of the implant.

### Comparison with SIMS literature on implants

In this study, it was discovered that more biomolecules were detected in the negative polarity, 247 metabolites, relative to the positive polarity which only detected 45 metabolites. This corroborates what was previously observed by Touboul *et al*. and Angerer *et al.* who detected more lipid species such as phosphoinositols, fatty acids and triglycerides in murine and human tissue sections in negative polarity relative to positive polarity analysis [54,55]. The complete list of MFP filtered and assigned peaks with reference to assignment databases are presented in **Table S1**-**S2**. The higher spectral and mass resolving power of cryo-OrbiSIMS and the GCIB primary beam has enabled us to detect and assign more biomolecular species on the surface of implants following extraction relative to the study conducted by Schreib *et al* [6]. Machine learning in combination with univariate statistical analysis has enabled us to narrow down the contribution of important of biomolecular species at the two time point of implant analysis. We showed that sugar-based molecules at 1 day were key features identified on the surface of the implant while lipid species such as fatty acids and urate were more important features at 28 days that indicate the presence of inflammation at the site of implant.

Previous work by Schreib *et al.* dissected and retrieved PDMS discs implanted in the intraperitoneal space of C57BL/6 mice, known to have a different immune profile compared to BALB/c mice. The BALB/c mouse strain, used in the current work, displays a more Th2 biased response in contrast to the C57bl/6 strain, that is more skewed towards a Th1 response.[56] For their PDMS implant, Schreib *et al.* reported a fibrotic collagen capsule of 33 ± 6 µm in thickness 28 days after implantation (n=3) into the intraperitoneal space. In the subcutaneous site from which our implants were retrieved, the collagen thickness was found to be thinner, 16 µm + 5 µm at 28 days implantation in BALB/c mice. These differences in collagen capsule thickness measured may be attributed to the different shape and size of implants used, different implantation site as well as the differences in fibrotic response expressed by the C57BL/6 mouse strain relative to the BALB/c mice used in the current work. These differences may result in the formation of a different FBR capsule with distinct structure along with different level of cell infiltration into the capsule relative to our current work. Schreib *et al.* reported no adhesion of the collagenous capsule to the explants, and a sparce cell distribution on the implant surface as visualised by microscopy. Our Masson’s trichrome staining allowed us to identify the presence of a cellular layer at the interface between implant site and the fibrous capsule. Due to complex immune and cellular cascade that take place between silicone-based implants and the host tissue, these cellular layers shown in **Figure 1(d)** may consist of a variety of cells ranging from monocytes, macrophage, T-cells and fibroblast [57].

Schreib *et al.* observed that the cells were surrounded by patches of molecules that contained the same secondary ion signatures as the plasma membrane. They suggested that this observation may be attributed to the cells leaving parts of their membrane on the surface of the disc upon contact and/or extracellular vesicles secreted from these sparsely distributed cells. With cryo-OrbiSIMS, we have been able to identify icosatetraenoic acid and glycerophosphoinositol PI (38:4) which are key marker for plasma membrane and endoplasmic reticulum [52,53]. The presence of this lipid may arise from host cell layer, as illustrated in **Figure 6(b)** depositing part of their membrane upon contact with the implant. It is worth noting the depth profile of the implant surface at 1 day showed that most of the nucleic acid bases are located deeper in the deposition away from the interface layer compared to 28 days. The presence of the nucleic acid may arise from the deposition of cellular residue such as through extracellular vesicle-associated DNA from the surrounding cells coming in close contact with the surface during the early phases of the implantation [58]. Also, the interface layer at 1 day consisted mostly of sugar isomers and its respective metabolites as shown in **Figure S8** and **Table S6**. The enrichment of sugars and metabolites at the interfacial layers may be attributed to increase in demand for energy particularly by macrophages and neutrophils at the implant site during the early inflammatory response [59]. This early inflammation is also a result of the disruption of cell structures due to the implantation procedure of the catheter into the subcutaneous pocket. We have highlighted that there is an increased in itaconate at the surface of the implant at 1 day *vs* 28 days. The depth profile data from **Figure S8** and **Table S6** showed that the molecule is spatially mostly located on the outermost layer of the deposit, near the interfacial layer. The presence of this anti-inflammatory molecule at the outermost layer of the deposits may serve as an to limit the inflammation at the site of implant and mitigate further damage to the periphery tissue.

Our GCIB/OrbiSIMS depth profile also showed that the outermost layer detected on the surface of the implant is enriched in lipids mostly of which are fatty acids, FA 16:0, FA 18:0, FA 18:1 and icosatetraenoic acid at 28 days. This observation is consistent with Schreib *et al.* who reported the enrichment of fatty acid on the surface of PDMS disk upon implantation into C57BL/6 mice at 1 day [6] using static LMIG ToF-SIMS analysis to analyse the surface lipid layer on their implant. However, due to limited spectral resolution of ToF-SIMS, they were unable to identify any molecular ions from other lipid and non-lipid species on the surface of their implant. In this study, with the aid of the Orbitrap analyser we also able to show that this interfacial layer is not only enriched with fatty acids but also contains nucleic acids bases such as guanine, adenine and cytosine. The DNA base adenine has been widely used as a nuclear cell marker for LMIG/ToF-SIMS and GCIB/Orbi-SIMS analysis [7,60]. The presence of uracil which is only present on the surface indicates the presence of RNA while the presence of thymine is an indicator that DNA is also present on the surface of the tubes in **Figure 6(a)**. In addition, due to inherent depth profiling capability of the Orbi-SIMS, we are also able to probe the presence of additional lipid species that may be present beneath this fatty acid enriched surface. Beneath this fatty acid enrich layer, we also showed the presence of other lipid species such as diacylglycerophosphoglycerol (PG 39:1;O), oxidised glycosphingolipids (SHexCer 41:5;O5), glycerophosphoinositols monophosphate (PIP 39:5) as well as the unsaturated linoleic acid (FA 18:2). This indeed builds upon the lipid hypothesis proposed Schreib *et al* that lipids deposition on the surface of biomaterial can play a critical role in biomaterial-induced foreign body reaction and fibrosis. However, this lipid deposition is not a homogeneous lipid layer but instead consist of layers of stratified lipid species. The ability to identify this stratified lipid layer was only achievable by the use of cryo GCIB/Orbi-SIMS analysis that enable the preservation of this lipid layer in a near native state during data acquisition.

### Conclusions

Cryo-OrbiSIMS using an argon GCIB to obtain high mass resolving power has been utilised to depth profile the biological deposit formed on the surface of silicone catheter implanted in mice at 1 day and 28 days. SIMS-MFP and logistic regression machine learning model have allowed the identification and characterisation of this complex biomolecular layer that consist of biomolecules of different composition at 1 and 28 days. Two distinct lipidic layers, with the outermost layer containing nucleic acid base and fatty acids has also been identified at the interface between implanted silicone catheter and host tissue providing molecular insight into the foreign body reaction at 28 days. This nucleic acid rich lipid interface forms the bridge that link between the biomaterials and the cellular layers that surrounds the biomaterials. It was shown that the elevated levels of urate, FA 14:0, FA 16:0 coupled with the presence of icosatetraenoic acid at the implant tissue interface provide indicator for increased inflammatory response at the implantation site. Such finding was corroborated by the observation of a highly inflamed subcutaneous tissue at 28 day which displayed the presence of collagen as shown *via* histological analysis. Overall, these findings show that cryo-GCIB/OrbiSIMS analytical workflow provides us with the necessary tools to deconvolute and understand the complex biointerface formed on silicone implant during an FBR response. Such methodology could be translatable to novel immune instructive surfaces enabling us to gain insights on the performance of biomaterials *in vivo*. Indeed, the expansion of this concept may provide an impetus to pave the way for advancements in developing novel immune-instructive materials with augmented functionality and biocompatibility.

## Conflict of interest and data availability

The authors declare no competing financial interest or personal relationships that could have appeared to influence the work reported in this paper.

## Dava Availability

The data that support the findings of this study are available under at the Nottingham Research Data Management Repository (http://doi.org/10.17639/nott.7533)

## Acknowledgements

The OrbiSIMS facility was funded by the EPSRC Strategic Equipment Grant ‘3D OrbiSIMS:Label free chemical imaging of materials, cells and tissues’ EP/P029868/1 and the work is also funded by the EPSRC Designing bio-instructive materials for translation-ready medical devices EP/X001156/1 2023-2027. Ian Gilmore is thanked for his many helpful discussions, including the suggestion to explore the use of a Volcano Plot for OrbiSIMS data.

