## Supplementary Information for "Molecular Biointerface Characterisation for an Implanted Medical Device Using Cryogenic Orbitrap Secondary Ion Mass Spectrometry (Cryo-OrbiSIMS)"

### Supporting Information

#### Data Analysis and Machine Learning

1. Data Pre-processing and Normalisation

The original dataset includes 60 PDMS samples (n=30 per time point) containing 282 attributes, considering only instances obtained under cryogenic temperature. OrbiSIMS spectra and profile that are collected after 24h and 28 days were included. These timeframes were defined as the dependent variables for the modelling exercise. The entire analytics was conducted using Helix[52], an open-source tool for data analysis and Machine Learning modelling and interpretation developed by our team. As the dataset is high dimensional and sparse, Least Absolute Shrinkage and Selection Operator (LASSO) was employed to eliminate less important variables before modelling. We also removed those variables with correlation above 0.95. This resulted in 54 independent variables to be used for modelling. Shapiro-Wilk and Kolgomorov-Smirnov tests reveal that most variables do not follow a normal distribution (**Table S3**).

2. Modelling

As the dataset is not normally distributed, we modelled the data using logistic regression and compared the results with non-linear Machine Learning approaches, namely, Random Forest, Support Vector Machine and XGBoost, using 5-fold cross-validation and 20% of the data for test. Hyperparameter search was conducted using grid search and results were collected for the models with best parameters. Improved results were obtained with the non-normalised data. Results are shown in **Table S4**. Logistic Regression and Random Forest produced the best classification results.

3. Post-Training Interpretation

To understand variable importance for logistic regression, we calculated the variable average beta coefficients across the 5 folds used for cross validation and their p-value to assess their significance, as shown in **Table S5**. A p-value smaller than 0.05 suggests statistical significance.

We use Shapley Additive explanation (SHAP) values to the logistic regression to further understand variable contributions to the outcome (**Table S4**). The variable 255.2328 appears to have the highest impact in the model, and its prevalence (high values represented as blue dots) affect more strongly data collected for 28 days. In the figure negative shap values indicate stronger correlation with 24h endpoint and positive correlate with 28 days data.

4. Machine Learning Interpretation

We utilised an ensemble post-training feature interpretation approach, implemented in Helix and detailed in the references provided in the comment, to better understand how variables influence both linear and non-linear model outcomes. Given the overall high accuracy achieved, we computed the mean and majority vote of the most important variables identified across all models (logistic regression, random forest, support vector machines, and XGBoost).

Overall, there was consensus regarding the most important variables, although differences in ranking emerged depending on the ensemble method employed. The **mean ensemble** quantifies variable importance across all machine learning models using two post-training methods: SHAP and Permutation Importance. In contrast, **majority voting** ranks variables based on their importance within each model and feature importance calculation, with the final ranking determined by the most frequently selected variables across methods.

**Table S1.** Tabulated output from the SIMS-MFP software showing the putative lipid assignments along with observed mass (m/z), mass deviation, chemical formula, molecular assignment and adduct. All assignments were compared against the Lipid Maps Structural Database®.

| mass | error | Formula | Assignment | Adduct |
|---|---|---|---|---|
| 549.4879 | 0.298 | $C_{35}H_{65}O_4^+$ | DG O-32:3/MG 32:3/MG O-32:4;O | [M+H] |
| 573.487 | -1.284 | $C_{37}H_{65}O_4^+$ | CE 10:0;O2/DG O-34:5/MG 34:5/MG O-34:6;O | [M+H] |
| 597.4869 | -1.399 | $C_{39}H_{65}O_4^+$ | CE 12:2;O2/DG O-36:7 | [M+H] |
| 551.5039 | 0.932 | $C_{35}H_{67}O_4^+$ | DG O-32:2/MG 32:2/MG O-32:3;O | [M+H] |
| 575.5039 | 0.893 | $C_{37}H_{67}O_4^+$ | DG O-34:4/MG 34:4/MG O-34:5;O | [M+H] |
| 599.5036 | 0.357 | $C_{39}H_{67}O_4^+$ | CE 12:1;O2/DG O-36:6 | [M+H] |
| 666.4845 | -1.802 | $C_{38}H_{69}NO_6P^+$ | CerP 38:5;O2/LPC O-30:6/LPE O-33:6 | [M+H] |
| 601.5191 | 0.106 | $C_{39}H_{69}O_4^+$ | CE 12:0;O2/DG O-36:5 | [M+H] |
| 621.4858 | 0.697 | $C_{34}H_{70}O_7P^+$ | LPA 31:0/LPA O-31:1;O/PA O-31:0 | [M+H] |
| 603.5353 | 1.017 | $C_{39}H_{71}O_4^+$ | DG O-36:4 | [M+H] |
| 623.5013 | 0.454 | $C_{34}H_{72}O_7P^+$ | LPA O-31:0;O | [M+H] |
| 725.5576 | 1.914 | $C_{42}H_{77}O_9^+$ | TG 39:2;O3 | [M+H] |
| 756.5527 | -1.429 | $C_{42}H_{79}NO_8P^+$ | CerP 42:4;O4/LPC 34:4;O/PC 34:3/PC O-34:4;O/ PE 37:3/PE O-37:4;O | [M+H] |
| 780.5524 | -1.769 | $C_{44}H_{79}NO_8P^+$ | CerP 44:6;O4/PC 36:5/PC O-36:6;O/PE 39:5/ PE O-39:6;O | [M+H] |
| 542.4906 | -0.275 | $C_{34}H_{65}NO_2Na^+$ | Cer 34:2;O/NAE 32:2 | [M+Na] |
| 621.4858 | 0.756 | $C_{39}H_{66}O_4Na^+$ | CE 12:1;O2/DG O-36:6 | [M+Na] |
| 623.5013 | 0.513 | $C_{39}H_{68}O_4Na^+$ | CE 12:0;O2/DG O-36:5 | [M+Na] |
| 577.517 | 0.641 | $C_{35}H_{70}O_4Na^+$ | DG O-32:0/MG 32:0/MG O-32:1;O | [M+Na] |
| 666.4845 | 1.808 | $C_{36}H_{70}NO_6PNa^+$ | CerP 36:2;O2/LPC O-28:3/LPE O-31:3 | [M+Na] |
| 625.5176 | 1.551 | $C_{39}H_{70}O_4Na^+$ | DG O-36:4 | [M+Na] |
| 725.5576 | 1.111 | $C_{39}H_{79}N_2O_6PNa^+$ | PE-Cer 37:1;O2/SM 34:1;O2 | [M+Na] |
| 773.5296 | -0.913 | $C_{40}H_{79}O_{10}PNa^+$ | BMP 34:0/LPG 34:1;O/PG 34:0/PG O-34:1;O | [M+Na] |
| 756.5527 | 1.751 | $C_{40}H_{80}NO_8PNa^+$ | CerP 40:1;O4/LPC 32:1;O/LPS O-34:1/PC 32:0/ PC O-32:1;O/PE 35:0/PE O-35:1;O | [M+Na] |
| 780.5524 | 1.313 | $C_{42}H_{80}NO_8PNa^+$ | CerP 42:3;O4/LPC 34:3;O/PC 34:2/PC O-34:3;O/ PE 37:2/PE O-37:3;O | [M+Na] |
| 690.4369 | -0.937 | $C_{34}H_{69}NO_8SK^+$ | NAT 32:0;O4 | [M+K] |
| 666.4845 | -1.975 | $C_{40}H_{69}NO_4K^+$ | CAR 33:5/Cer 40:6;O3 | [M+K] |
| 621.4858 | 0.512 | $C_{36}H_{70}O_5K^+$ | DG 33:0/DG O-33:1;O/MG 33:1;O/TG O-33:0 | [M+K] |
| 623.5013 | 0.269 | $C_{36}H_{72}O_5K^+$ | DG O-33:0;O/MG 33:0;O | [M+K] |
| 756.5527 | -1.581 | $C_{44}H_{79}NO_6K^+$ | ACer 44:4;O4/Cer 44:5;O5 | [M+K] |

| | | | | |
|---|---|---|---|---|
| 780.5524 | -1.917 | $C_{46}H_{79}NO_6K^+$ | ACer 46:6;O4 | [M+K] |

| mass | error | Formula | Assignment | Adduct |
|---|---|---|---|---|
| 227.2016 | -0.237 | $C_{14}H_{27}O_2^-$ | FA 14:0 | [M-H] |
| 299.2014 | -0.849 | $C_{20}H_{27}O_2^-$ | ST 20:3;O2 | [M-H] |
| 253.2174 | 0.380 | $C_{16}H_{29}O_2^-$ | FA 16:1 | [M-H] |
| 301.2172 | -0.345 | $C_{20}H_{29}O_2^-$ | FA 20:5/ST 20:2;O2 | [M-H] |
| 255.2328 | -0.603 | $C_{16}H_{31}O_2^-$ | FA 16:0 | [M-H] |
| 279.2326 | -1.267 | $C_{18}H_{31}O_2^-$ | FA 18:2 | [M-H] |
| 303.2327 | -0.837 | $C_{20}H_{31}O_2^-$ | FA 20:4/ST 20:1;O2 | [M-H] |
| 327.2326 | -1.081 | $C_{22}H_{31}O_2^-$ | FA 22:6/ST 22:3;O2 | [M-H] |
| 269.2488 | 0.728 | $C_{17}H_{33}O_2^-$ | FA 17:0 | [M-H] |
| 281.2486 | -0.014 | $C_{18}H_{33}O_2^-$ | FA 18:1 | [M-H] |
| 305.2483 | -0.996 | $C_{20}H_{33}O_2^-$ | FA 20:3/ST 20:0;O2 | [M-H] |
| 329.2484 | -0.619 | $C_{22}H_{33}O_2^-$ | FA 22:5/ST 22:2;O2 | [M-H] |
| 283.2642 | -0.190 | $C_{18}H_{35}O_2^-$ | FA 18:0 | [M-H] |
| 331.2639 | -1.068 | $C_{22}H_{35}O_2^-$ | FA 22:4/ST 22:1;O2 | [M-H] |
| 391.2251 | -1.022 | $C_{19}H_{36}O_6P^-$ | LPA O-16:2 | [M-H] |
| 420.2601 | -0.459 | $C_{20}H_{38}NO_8^-$ | CAR 13:0;O4 | [M-H] |
| 417.2411 | -0.120 | $C_{21}H_{38}O_6P^-$ | LPA O-18:3 | [M-H] |
| 359.2954 | -0.428 | $C_{24}H_{39}O_2^-$ | FA 24:4/ST 24:1;O2 | [M-H] |
| 419.2574 | 1.431 | $C_{21}H_{40}O_6P^-$ | LPA O-18:2 | [M-H] |
| 437.2675 | 0.309 | $C_{21}H_{42}O_7P^-$ | LPA 18:0/LPA O-18:1;O | [M-H] |
| 436.2835 | 0.346 | $C_{21}H_{43}NO_6P^-$ | LPC O-13:1/LPE O-16:1 | [M-H] |
| 462.2992 | 0.435 | $C_{23}H_{45}NO_6P^-$ | LPC O-15:2/LPE O-18:2 | [M-H] |
| 464.3142 | -0.967 | $C_{23}H_{47}NO_6P^-$ | LPC O-15:1/LPE O-18:1 | [M-H] |
| 480.3098 | 0.492 | $C_{23}H_{47}NO_7P^-$ | LPC 15:0/LPC O-15:1;O/LPE 18:0/LPE O-18:1;O | [M-H] |
| 588.3738 | 1.667 | $C_{34}H_{54}NO_5S^-$ | NAT 32:7;O | [M-H] |
| 616.4721 | 1.543 | $C_{34}H_{67}NO_6P^-$ | CerP 34:1;O2/LPC O-26:2/LPE O-29:2 | [M-H] |
| 905.3844 | -1.658 | $C_{42}H_{67}O_{17}P_2^-$ | PIP 33:7;O | [M-H] |
| 642.4864 | -0.621 | $C_{36}H_{69}NO_6P^-$ | CerP 36:2;O2/LPC O-28:3/LPE O-31:3 | [M-H] |
| 722.5129 | -0.158 | $C_{41}H_{73}NO_7P^-$ | CerP 41:5;O3/LPC 33:5/LPC O-33:6;O/PC O-33:5/PE O-36:5 | [M-H] |
| 746.514 | 1.321 | $C_{43}H_{73}NO_7P^-$ | PC O-35:7/PE O-38:7 | [M-H] |
| 748.5296 | 1.251 | $C_{43}H_{75}NO_7P^-$ | CerP 43:6;O3/PC O-35:6/PE O-38:6 | [M-H] |

| | | | | |
|---|---|---|---|---|
| 723.517 | -1.603 | $C_{38}H_{76}O_{10}P^-$ | LPG 32:0;O/PG O-32:0;O | [M-H] |
| 687.5443 | -0.506 | $C_{38}H_{76}N_2O_6P^-$ | PE-Cer 36:1;O2/SM 33:1;O2 | [M-H] |
| 747.517 | -1.552 | $C_{40}H_{76}O_{10}P^-$ | BMP 34:1/LPG 34:2;O/PG 34:1/PG O-34:2;O | [M-H] |
| 858.5235 | 1.691 | $C_{44}H_{76}NO_{15}^-$ | Hex2Cer 32:4;O4 | [M-H] |
| 718.5382 | -1.432 | $C_{39}H_{77}NO_8P^-$ | CerP 39:1;O4/LPC 31:1;O/LPE 34:1;O/LPS O-33:1/PC 31:0/PC O-31:1;O/PE 34:0/PE O-34:1;O | [M-H] |
| 742.539 | -0.308 | $C_{41}H_{77}NO_8P^-$ | CerP 41:3;O4/LPC 33:3;O/PC 33:2/PC O-33:3;O/PE 36:2/PE O-36:3;O | [M-H] |
| 750.5438 | -0.685 | $C_{43}H_{77}NO_7P^-$ | CerP 43:5;O3/PC O-35:5/PE O-38:5 | [M-H] |
| 766.5398 | 0.745 | $C_{43}H_{77}NO_8P^-$ | CerP 43:5;O4/PC 35:4/PC O-35:5;O/PE 38:4/PE O-38:5;O | [M-H] |
| 994.4794 | 1.198 | $C_{46}H_{77}NO_{20}P^-$ | MIPC 34:6;O6 | [M-H] |
| 857.5185 | -0.064 | $C_{45}H_{78}O_{13}P^-$ | PI 36:4/PI O-36:5;O | [M-H] |
| 744.555 | 0.163 | $C_{41}H_{79}NO_8P^-$ | CerP 41:2;O4/LPC 33:2;O/PC 33:1/PC O-33:2;O/PE 36:1/PE O-36:2;O | [M-H] |
| 886.5531 | -0.280 | $C_{46}H_{80}NO_{15}^-$ | Hex2Cer 34:4;O4 | [M-H] |
| 883.5344 | 0.221 | $C_{47}H_{80}O_{13}P^-$ | PI 38:5/PI O-38:6;O | [M-H] |
| 778.5754 | -0.275 | $C_{45}H_{81}NO_7P^-$ | CerP 45:5;O3/PC O-37:5/PE O-40:5 | [M-H] |
| 794.5717 | 1.474 | $C_{45}H_{81}NO_8P^-$ | CerP 45:5;O4/PC 37:4/PC O-37:5;O/PE 40:4/PE O-40:5;O | [M-H] |
| 861.5491 | -0.876 | $C_{45}H_{82}O_{13}P^-$ | PI 36:2/PI O-36:3;O | [M-H] |
| 885.549 | -0.965 | $C_{47}H_{82}O_{13}P^-$ | PI 38:4/PI O-38:5;O | [M-H] |
| 814.5983 | 1.910 | $C_{45}H_{85}NO_9P^-$ | CerP 45:3;O5/PC 37:2;O/PE 40:2;O/PS O-39:2 | [M-H] |
| 910.5677 | 1.610 | $C_{45}H_{85}NO_{15}P^-$ | PI-Cer 39:2;O6 | [M-H] |
| 833.5926 | 1.530 | $C_{45}H_{86}O_{11}P^-$ | PG 39:1;O | [M-H] |
| 813.6008 | -0.854 | $C_{46}H_{86}O_9P^-$ | PA 43:2;O/PG O-40:3 | [M-H] |
| 711.6513 | 0.683 | $C_{44}H_{87}O_6^-$ | DG O-41:0;O2 | [M-H] |
| 795.6392 | 0.820 | $C_{46}H_{88}N_2O_6P^-$ | PE-Cer 44:3;O2/SM 41:3;O2 | [M-H] |
| 810.6464 | -0.071 | $C_{47}H_{88}NO_9^-$ | HexCer 41:2;O3 | [M-H] |
| 790.6345 | 1.734 | $C_{44}H_{89}NO_8P^-$ | CerP 44:0;O4/PC O-36:0;O/PE O-39:0;O | [M-H] |
| 797.6543 | 0.128 | $C_{46}H_{90}N_2O_6P^-$ | PE-Cer 44:2;O2/SM 41:2;O2 | [M-H] |
| 952.5239 | 1.122 | $C_{47}H_{83}NO_{14}SCl^-$ | SHexCer 41:5;O5 | [M+Cl] |
| 1013.492 | -1.049 | $C_{48}H_{84}O_{16}P_2Cl^-$ | PIP 39:5 | [M+Cl] |
| 790.6345 | 1.512 | $C_{45}H_{89}NO_7Cl^-$ | ACer 45:0;O5/Cer 45:1;O6 | [M+Cl] |

**Table S2.** Tabulated output from the SIMS-MFP software showing the putative biomolecular assignments along with observed mass (m/z), mass deviation, chemical formula, molecular assignment and adduct. All assignments were compared against the Human Metabolome Database®.

| mass | error | formula | Assignment | Adduct |
|---|---|---|---|---|
| 206.9961 | 1.512 | $C_6H_7O_6S$ | Pyrogallol-2-O-sulphate | [M+H] |
| 130.0653 | 1.345 | $C_9H_8N$ | leucoline | [M+H] |
| 173.0269 | 1.202 | $C_7H_9O_3S$ | p-Toluenesulfonic acid | [M+H] |
| 168.0588 | -1.155 | $C_7H_{10}N_3S$ | 6-Methyl-7,8-dihydroimidazo[1,5-c]pyrimidine-5(6H)-thione | [M+H] |
| 91.0574 | -2.165 | $C_4H_{11}S$ | butanethiol | [M+H] |
| 107.0523 | -1.981 | $C_4H_{11}OS$ | methionol | [M+H] |
| 179.0553 | 1.586 | $C_6H_{11}O_6$ | L-Gulono-14-lactone/ D-Glucono-14-lactone | [M+H] |
| 86.0966 | 2.031 | $C_5H_{12}N$ | Piperidine | [M+H] |
| 109.1013 | 1.136 | $C_8H_{13}$ | (Z)-1,3-Octadiene | [M+H] |
| 265.0378 | 0.558 | $C_9H_{13}O_7S$ | 3-Methoxy-4-Hydroxyphenylglycolsulfate | [M+H] |
| 184.0736 | 1.515 | $C_5H_{15}NO_4P$ | Choline phosphate | [M+H] |
| 250.0085 | -0.840 | $C_5H_{10}NO_7PNa$ | Î³-Glutamyl phosphate | [M+Na] |
| 270.0142 | 1.500 | $C_8H_{10}NO_6PNa$ | Pyridoxal phosphate | [M+Na] |
| 184.0736 | 1.716 | $C_{10}H_{11}NONa$ | Tryptophol | [M+Na] |
| 759.5121 | -1.189 | $C_{41}H_{72}N_2O_9Na$ | oceanalin A | [M+Na] |
| 773.5296 | -0.913 | $C_{40}H_{79}O_{0}PNa$ | 1-hexadecanoyl-2-octadecanoyl-sn-glycero-3-phospho-(1 -sn-glycerol) | [M+Na] |
| 124.9997 | -1.897 | $C_4H_6O_2K$ | 3-Butenoic acid/ Biacetyl/ butyrolactone/ oxolan-3-one | [M+K] |
| 157.016 | -1.625 | $C_7H_6N_2K$ | pyrrolo[1,2-a]pyrazine | [M+K] |
| 216.99 | 1.073 | $C_9H_6O_4K$ | 6,7-Dihydroxycoumarin | [M+K] |
| 231.0055 | 0.359 | $C_{10}H_8O_4K$ | Scopoletin | [M+K] |
| 211.0271 | 1.326 | $C_{10}H_8N_2OK$ | 4-Hydroxy-1H-indole-3-acetonitrile | [M+K] |
| 192.9756 | 1.145 | $C_4H_{10}O_2S_2K$ | 2-hydroxyethyldisulfide | [M+K] |
| 237.9879 | 0.705 | $C_4H_{10}NO_6PK$ | (S)-2-Amino-3-hydroxybutanoic acid 3-phosphate | [M+K] |
| 265.9829 | 0.951 | $C_5H_{10}NO_7PK$ | Î³-Glutamyl phosphate | [M+K] |
| 249.016 | 0.072 | $C_{10}H_{10}O_5K$ | 3-(4-Hydroxy-3-methoxyphenyl)-2-oxiranecarboxylic acid | [M+K] |
| 228.0385 | 1.692 | $C_6H_{11}N_3O_4K$ | gly-asn | [M+K] |
| 227.0427 | -0.727 | $C_7H_{12}N_2O_4K$ | N-Acetylglutamine | [M+K] |
| 184.0736 | 0.890 | $C_7H_{15}NO_2K$ | 4-Trimethylammoniobutanoate/4-Trimethylammoniobutanoate/ g-Butyrobetaine | [M+K] |

| mass | error | formula | Assignment | Adduct |
|---|---|---|---|---|
| 82.9958 | -3.553 | $C_4H_3S^-$ | Thiophene | [M-H] |
| 83.0136 | -3.059 | $C_4H_3O_2^-$ | 2[5H]-furanone | [M-H] |
| 99.0086 | -1.706 | $C_4H_3O_3^-$ | Succinic anhydride | [M-H] |
| 111.0199 | -0.918 | $C_4H_3N_2O_2^-$ | Uracil/ 4-CARBOXYPYRAZOLE | [M-H] |
| 111.0087 | -0.621 | $C_5H_3O_3^-$ | 2-Furoic acid | [M-H] |
| 127.0039 | 1.702 | $C_5H_3O_4^-$ | 5-Hydroxy-2-furoic acid | [M-H] |
| 123.0198 | -1.641 | $C_5H_3N_2O_2^-$ | Pyrazinoate/ Pyrazinoic acid | [M-H] |
| 135.031 | -1.740 | $C_5H_3N_4O^-$ | Hypoxanthine | [M-H] |
| 167.0208 | -1.586 | $C_5H_3N_4O_3^-$ | Urate/ Uric Acid | [M-H] |
| 110.036 | 0.128 | $C_4H_4N_3O^-$ | Cytosine | [M-H] |
| 126.0308 | -0.801 | $C_4H_4N_3O_2^-$ | (3r,4s)-Tofacitinib | [M-H] |
| 94.0294 | -4.657 | $C_5H_4NO^-$ | 2-pyridone | [M-H] |
| 126.0018 | -0.864 | $C_5H_4NOS^-$ | 2-Acetylthiazole | [M-H] |
| 110.0246 | -1.390 | $C_5H_4NO_2^-$ | Pyrrole 2-carboxylate | [M-H] |
| 126.0196 | -0.539 | $C_5H_4NO_3^-$ | 4-Hydroxy-1H-pyrrole-2-carboxylic acid | [M-H] |
| 134.047 | -1.633 | $C_5H_4N_5^-$ | Adenine | [M-H] |
| 150.042 | -0.892 | $C_5H_4N_5O_1^-$ | Guanine | [M-H] |
| 122.0246 | -1.253 | $C_6H_4NO_2^-$ | Picolinic acid/ Isonicotinic acid | [M-H] |
| 138.0195 | -1.216 | $C_6H_4NO_3^-$ | 4-Nitrophenol | [M-H] |
| 146.0245 | -1.732 | $C_8H_4NO_2^-$ | Indole-5,6-quinone/ Isatin | [M-H] |
| 85.0292 | -3.574 | $C_4H_5O_2^-$ | 3-Butenoic acid/ Biacetyl/ butyrolactone/ oxolan-3-one | [M-H] |
| 133.0145 | 1.888 | $C_4H_5O_5^-$ | Malate/ (±)-Malic Acid | [M-H] |
| 149.0093 | 0.913 | $C_4H_5O_6^-$ | (.+/-.)-Tartaric acid | [M-H] |
| 113.0356 | -0.459 | $C_4H_5N_2O_2^-$ | Dihydrouracil/ Dioxy-creatinine | [M-H] |
| 81.0342 | -4.799 | $C_5H_5O^-$ | 2-Methylfuran | [M-H] |
| 113.0244 | -0.167 | $C_5H_5O_3^-$ | 3-Oxo-4-pentenoic acid/ 4-Hydroxy-5-methylfuran-3(2H)-one | [M-H] |
| 129.0192 | -1.038 | $C_5H_5O_4^-$ | Itaconate/ Mesaconate/ (Z)-glutaconic acid/ 4,5-dioxovaleric acid | [M-H] |
| 93.0456 | -2.385 | $C_5H_5N_2^-$ | Methylpyrazine | [M-H] |
| 125.0356 | -0.415 | $C_5H_5N_2O_2^-$ | Thymine/ Imidazole-4-acetate | [M-H] |
| 125.0243 | -0.951 | $C_6H_5O_3^-$ | (3E)-2,5-Dioxo-3-hexenal/ Pyrogallol | [M-H] |
| 172.9913 | -0.606 | $C_6H_5O_4S^-$ | Phenol sulfate | [M-H] |
| 173.0094 | 1.365 | $C_6H_5O_6^-$ | trans-Aconitate/ L-Dehydroascorbate/ (E)-Aconitic Acid | [M-H] |

| | | | | |
|---|---|---|---|---|
| 204.9813 | 0.318 | $C_6H_5O_6S^-$ | Pyrogallol-2-O-sulphate | [M-H] |
| 121.0406 | -1.131 | $C_6H_5N_2O^-$ | Nicotinamide | [M-H] |
| 137.0355 | -1.108 | $C_6H_5N_2O_2^-$ | Urocanic acid | [M-H] |
| 121.0294 | -0.858 | $C_7H_5O_2^-$ | Benzoate/ 4-Hydroxybenzaldehyde/ Benzoic acid | [M-H] |
| 117.0457 | -1.041 | $C_7H_5N_2^-$ | pyrrolo[1,2-a]pyrazine | [M-H] |
| 117.0345 | -0.760 | $C_8H_5O^-$ | 2,4,6-Octatriyn-1-ol/ Coumarone | [M-H] |
| 145.0293 | -1.406 | $C_9H_5O_2^-$ | Coumarin | [M-H] |
| 84.0452 | -3.426 | $C_4H_6NO^-$ | 2-Pyrrolidone | [M-H] |
| 116.0353 | -0.154 | $C_4H_6NO_3^-$ | Aceturic acid | [M-H] |
| 112.0515 | -1.213 | $C_4H_6N_3O^-$ | Creatinine | [M-H] |
| 80.0502 | -4.658 | $C_5H_6N^-$ | methyl pyrrole | [M-H] |
| 112.0405 | 0.867 | $C_5H_6NO_2^-$ | 1-pyrroline-5-carboxylic acid | [M-H] |
| 128.0352 | -0.921 | $C_5H_6NO_3^-$ | 5-Oxoproline/ Pyroglutamic acid | [M-H] |
| 108.0567 | -0.193 | $C_5H_6N_3^-$ | 2-Amino-4-methylpyrimidine | [M-H] |
| 124.0515 | -1.096 | $C_5H_6N_3O^-$ | 5-Methylcytosine | [M-H] |
| 92.0503 | -2.965 | $C_6H_6N^-$ | Aniline | [M-H] |
| 108.0454 | -0.814 | $C_6H_6NO^-$ | 4-Aminophenol | [M-H] |
| 140.0352 | -0.842 | $C_6H_6NO_3^-$ | 2-Aminomuconic acid semialdehyde | [M-H] |
| 136.0514 | -1.734 | $C_6H_6N_3O^-$ | Isoniazid | [M-H] |
| 136.0402 | -1.491 | $C_7H_6NO_2^-$ | 4-Aminobenzoate/ 4-Aminobenzoic acid | [M-H] |
| 152.0352 | -0.775 | $C_7H_6NO_3^-$ | Mesalazine | [M-H] |
| 116.0505 | -0.628 | $C_8H_6N^-$ | Indole | [M-H] |
| 164.0352 | -0.719 | $C_8H_6NO_3^-$ | 3-Hydroxy-2H-indole-5,6(3H,7H)-dione/ Coixol | [M-H] |
| 160.0403 | -0.643 | $C_9H_6NO_2^-$ | 3-Carboxyindole | [M-H] |
| 119.0171 | -1.049 | $C_4H_7O_2S^-$ | 4-Thiapentanoic acid | [M-H] |
| 99.0563 | -0.877 | $C_4H_7N_2O^-$ | NPYR | [M-H] |
| 115.0512 | -0.886 | $C_4H_7N_2O_2^-$ | 1-Nitroso-3-pyrrolidinol/ Succinamide | [M-H] |
| 83.0499 | -4.081 | $C_5H_7O^-$ | Cyclopentanone | [M-H] |
| 115.0401 | 0.270 | $C_5H_7O_3^-$ | 3-Methyl-2-oxobutanoic acid/ Levulinic acid | [M-H] |
| 127.0512 | -0.802 | $C_5H_7N_2O_2^-$ | 5-Nitrilonorvaline | [M-H] |
| 111.045 | -1.386 | $C_6H_7O_2^-$ | Sorbic acid | [M-H] |
| 159.0301 | 1.265 | $C_6H_7O_5^-$ | 2-Oxoadipate | [M-H] |
| 107.0614 | -0.672 | $C_6H_7N_2^-$ | Phenylhydrazine/ 2,5-Dimethylpyrazine | [M-H] |
| 123.0563 | -0.706 | $C_6H_7N_2O^-$ | 2-Methoxy-3-methylpyrazine | [M-H] |
| 139.0512 | -0.733 | $C_6H_7N_2O_2^-$ | 1,3-dimethyluracil | [M-H] |

| | | | | |
|---|---|---|---|---|
| 155.046 | -1.399 | $C_6H_7N_2O_3^-$ | pentoxyl | [M-H] |
| 91.0551 | -2.459 | $C_7H_7^-$ | Toluene | [M-H] |
| 107.0501 | -1.298 | $C_7H_7O^-$ | Anisole | [M-H] |
| 123.045 | -1.251 | $C_7H_7O_2^-$ | Guaiacol | [M-H] |
| 171.0118 | -1.988 | $C_7H_7O_3S^-$ | p-Toluenesulfonic acid | [M-H] |
| 203.0196 | -0.635 | $C_7H_7O_7^-$ | 2,6-Anhydro-6-carboxy-3-deoxyhex-2-enonic acid | [M-H] |
| 119.0614 | -0.604 | $C_7H_7N_2^-$ | 2-Methyl-5-vinylpyrazine | [M-H] |
| 135.0563 | -0.643 | $C_7H_7N_2O^-$ | Nicotinyl methylamide | [M-H] |
| 167.046 | -1.298 | $C_7H_7N_2O_3^-$ | N-(1H-Pyrrol-2-ylcarbonyl)glycine | [M-H] |
| 119.0501 | -1.167 | $C_8H_7O^-$ | 7-Octene-3,5-diyn-1-ol | [M-H] |
| 135.045 | -1.140 | $C_8H_7O_2^-$ | 3-(5-methyl-2-furyl)prop-2-enal | [M-H] |
| 183.0119 | -1.311 | $C_8H_7O_3S^-$ | 4-styrenesulfonic acid | [M-H] |
| 131.0613 | -1.312 | $C_8H_7N_2^-$ | 1-Methylpyrrolo[1,2-a]pyrazine | [M-H] |
| 147.0449 | -1.727 | $C_9H_7O_2^-$ | 2-Chromanone/ Cinnamic acid | [M-H] |
| 143.0502 | -0.272 | $C_{10}H_7O^-$ | 1-Naphthol | [M-H] |
| 171.0563 | -0.508 | $C_{10}H_7N_2O^-$ | 4-Hydroxy-1H-indole-3-acetonitrile | [M-H] |
| 102.056 | -0.518 | $C_4H_8NO_2^-$ | 4-Aminobutanoate/ gamma-Aminobutyric acid | [M-H] |
| 130.0621 | -0.776 | $C_4H_8N_3O_2^-$ | Creatine | [M-H] |
| 114.056 | -0.464 | $C_5H_8NO_2^-$ | L-Proline | [M-H] |
| 130.0508 | -1.291 | $C_5H_8NO_3^-$ | 5-Aminolevulinate/ Ac-Ala-OH/ Aminolevulinic acid/ L-Hydroxyproline | [M-H] |
| 110.0723 | -0.644 | $C_5H_8N_3^-$ | 1H-Imidazole-4-ethanamine/2-Aminohistamine/ Histamine | [M-H] |
| 110.0611 | -0.344 | $C_6H_8NO^-$ | 2-Acetyl-1-pyrroline | [M-H] |
| 126.0559 | -1.213 | $C_6H_8NO_2^-$ | D-1-Piperideine-2-carboxylic acid | [M-H] |
| 142.0508 | -1.182 | $C_6H_8NO_3^-$ | 6-Oxo-pipecolinic acid/ trimethadione | [M-H] |
| 138.0671 | -1.346 | $C_6H_8N_3O^-$ | 2-Amino-3-(1H-imidazol-4-yl)propanal | [M-H] |
| 154.062 | -1.304 | $C_6H_8N_3O_2^-$ | L-Histidine | [M-H] |
| 170.057 | -0.682 | $C_6H_8N_3O_3^-$ | Metronidazole | [M-H] |
| 106.0662 | -0.216 | $C_7H_8N^-$ | Benzylamine | [M-H] |
| 138.0559 | -1.108 | $C_7H_8NO_2^-$ | Ethyl 2-pyrrolecarboxylate | [M-H] |
| 150.067 | -1.905 | $C_7H_8N_3O^-$ | 4-phenylsemicarbazide | [M-H] |
| 134.061 | -1.029 | $C_8H_8NO^-$ | 2-Aminoacetophenone/ Acetanilide | [M-H] |
| 130.0661 | -0.945 | $C_9H_8N^-$ | Skatole | [M-H] |
| 146.061 | -0.944 | $C_9H_8NO^-$ | Indole-3-carbidol | [M-H] |

| | | | | |
|---|---|---|---|---|
| 162.0559 | -0.944 | $C_9H_8NO_2^-$ | Î³-Oxo-3-pyridinebutanal | [M-H] |
| 158.061 | -0.872 | $C_{10}H_8NO^-$ | Indole-3-acetaldehyde/ N-hydroxy-1-naphthylamine | [M-H] |
| 186.056 | -0.284 | $C_{11}H_8NO_2^-$ | Indoleacrylic acid | [M-H] |
| 89.0426 | -4.996 | $C_4H_9S^-$ | butanethiol | [M-H] |
| 105.0378 | -1.522 | $C_4H_9OS^-$ | methionol | [M-H] |
| 149.0453 | -1.670 | $C_5H_9O_5^-$ | D-Ribose/ D-Arabinose/ 2-deoxy-D-ribonic acid/ D-(+)-Xylose/ D-Ribofuranose/ D-Xylopyranose | [M-H] |
| 165.0407 | 1.431 | $C_5H_9O_6^-$ | L-Xylonic acid | [M-H] |
| 113.072 | -0.326 | $C_5H_9N_2O^-$ | 3-Amino-2-piperidinone | [M-H] |
| 145.0617 | -1.151 | $C_5H_9N_2O_3^-$ | L-Glutamine | [M-H] |
| 161.0458 | 1.559 | $C_6H_9O_5^-$ | 2-Hydroxyadipic acid/ Diethylpyrocarbonate/ Meglutol | [M-H] |
| 177.0402 | -1.491 | $C_6H_9O_6^-$ | L-Gulono-14-lactone/ D-Glucono-14-lactone | [M-H] |
| 121.077 | -1.007 | $C_7H_9N_2^-$ | 2,4-Diaminotoluene | [M-H] |
| 137.0719 | -0.999 | $C_7H_9N_2O^-$ | 2-Ethyl-3-methoxypyrazine | [M-H] |
| 121.0658 | -0.734 | $C_8H_9O^-$ | Phenylethyl alcohol | [M-H] |
| 133.077 | -0.916 | $C_8H_9N_2^-$ | Cyclohexapyrazine | [M-H] |
| 133.0657 | -1.420 | $C_9H_9O^-$ | 3-Phenylpropanal/ Cinnamyl alcohol | [M-H] |
| 161.0719 | -0.850 | $C_9H_9N_2O^-$ | norcotinine | [M-H] |
| 193.0616 | -1.382 | $C_9H_9N_2O_3^-$ | 4-Aminohippuricacid | [M-H] |
| 116.0716 | -0.887 | $C_5H_{10}NO_2^-$ | L-Valine/ Betaine | [M-H] |
| 144.0776 | -1.741 | $C_5H_{10}N_3O_2^-$ | g-Guanidinobutyrate | [M-H] |
| 241.0115 | -1.576 | $C_6H_{10}O_8P^-$ | 1D-myo-Inositol 1,2-cyclic phosphate (Inositol cyclic phosphate) | [M-H] |
| 140.0829 | -0.256 | $C_6H_{10}N_3O^-$ | histidinol | [M-H] |
| 124.0766 | -1.514 | $C_7H_{10}NO^-$ | 2-Ethyl-4,5-dimethylthiazole | [M-H] |
| 140.0716 | -0.735 | $C_7H_{10}NO_2^-$ | Ethosuximide | [M-H] |
| 152.0829 | -0.236 | $C_7H_{10}N_3O^-$ | Acetylhistamine | [M-H] |
| 168.0777 | -0.898 | $C_7H_{10}N_3O_2^-$ | N(pi)-Methyl-L-histidine | [M-H] |
| 184.0726 | -0.901 | $C_7H_{10}N_3O_3^-$ | N-(1-Methyl-4-oxo-4,5-dihydro-1H-imidazol-2-yl)alanine | [M-H] |
| 136.0766 | -1.381 | $C_8H_{10}NO^-$ | Tyramine | [M-H] |
| 152.0715 | -1.334 | $C_8H_{10}NO_2^-$ | dopamine | [M-H] |
| 196.073 | 1.194 | $C_8H_{10}N_3O_3^-$ | N-Acetyl-DL-histidine | [M-H] |
| 148.0766 | -1.269 | $C_9H_{10}NO^-$ | 5-acetyl-2,3-dihydro-1H-pyrrolizine | [M-H] |
| 164.0715 | -1.237 | $C_9H_{10}NO_2^-$ | L-Phenylalanine | [M-H] |
| 176.0827 | -1.340 | $C_9H_{10}N_3O^-$ | NNN | [M-H] |

| | | | | |
|---|---|---|---|---|
| 160.0765 | -1.799 | $C_{10}H_{10}NO^-$ | Tryptophol | [M-H] |
| 172.0766 | -1.092 | $C_{11}H_{10}NO^-$ | pyroquilon | [M-H] |
| 188.0716 | -0.547 | $C_{11}H_{10}NO_2^-$ | phensuximide | [M-H] |
| 168.0431 | -0.112 | $C_4H_{11}NO_4P^-$ | P-DMEA | [M-H] |
| 131.0824 | -1.540 | $C_5H_{11}N_2O_2^-$ | L-Ornithine | [M-H] |
| 179.0558 | -1.753 | $C_6H_{11}O_6^-$ | myo-Inositol/ Fructose/ Glucose/ Galactose/ beta-D-Galactopyranose | [M-H] |
| 135.0813 | -1.769 | $C_9H_{11}O^-$ | Nonatrienal/ Mesitol | [M-H] |
| 179.0825 | -0.569 | $C_9H_{11}N_2O_2^-$ | (4-Ethoxyphenyl)urea | [M-H] |
| 195.0773 | -1.112 | $C_9H_{11}N_2O_3^-$ | 5-Nitro-2-propoxyaniline | [M-H] |
| 147.0813 | -1.624 | $C_{10}H_{11}O^-$ | Estragole | [M-H] |
| 175.0875 | -1.067 | $C_{10}H_{11}N_2O^-$ | Serotonin | [M-H] |
| 191.0825 | -0.533 | $C_{10}H_{11}N_2O_2^-$ | cotinine N-oxide/ Hydroxycotinine | [M-H] |
| 207.0773 | -1.047 | $C_{10}H_{11}N_2O_3^-$ | L-Kynurenine | [M-H] |
| 203.0825 | -0.502 | $C_{11}H_{11}N_2O_2^-$ | L-Tryptophan | [M-H] |
| 130.0985 | -0.660 | $C_5H_{12}N_3O^-$ | 1-(4-Aminobutyl)urea | [M-H] |
| 259.0224 | -0.173 | $C_6 H_{12}O_9P^-$ | D-Fructose 6-phosphate/ D-glucose 6-phosphate/ D-glucopyranose 6-phosphate | [M-H] |
| 130.0872 | -1.175 | $C_6 H_{12}NO_2^-$ | L-Norleucine/ L-Leucine/ L-Isoleucine/ L-Alloisoleucine | [M-H] |
| 126.0923 | -1.094 | $C_7H_{12}NO^-$ | 8-Azabicyclo[3.2.1]octan-3-ol | [M-H] |
| 138.0923 | -0.999 | $C_8H_{12}NO^-$ | Norpseudopelletierine | [M-H] |
| 154.0872 | -0.992 | $C_8H_{12}NO_2^-$ | arecoline | [M-H] |
| 166.0871 | -1.523 | $C_9H_{12}NO_2^-$ | 3-Methoxytyramine | [M-H] |
| 194.0934 | -0.520 | $C_9H_{12}N_3O_2^-$ | 4-(Nitrosoamino)-1-(3-pyridinyl)-1-butanol | [M-H] |
| 210.0883 | -0.552 | $C_9H_{12}N_3O_3^-$ | Zalcitabine | [M-H] |
| 129.1032 | -1.060 | $C_6H_{13}N_2O^-$ | N-Acetylputrescine/ N-Acetylputrescine | [M-H] |
| 145.0981 | -1.047 | $C_6H_{13}N_2O_2^-$ | L-Lysine | [M-H] |
| 169.0981 | -0.898 | $C_8H_{13}N_2O_2^-$ | R-(+)-Etiracetam | [M-H] |
| 165.1032 | -0.829 | $C_9H_{13}N_2O^-$ | 2-Isobutyl-3-methoxypyrazine | [M-H] |
| 193.0981 | -0.787 | $C_{10}H_{13}N_2O_2^-$ | 6-hydroxypseudooxynicotine | [M-H] |
| 209.093 | -0.798 | $C_{10}H_{13}N_2O_3^-$ | Aprobarbital | [M-H] |
| 189.103 | -1.782 | $C_{11}H_{13}N_2O^-$ | 5-Methoxytryptamine | [M-H] |
| 140.1079 | -1.341 | $C_8H_{14}NO^-$ | Isopelletierine | [M-H] |
| 196.1089 | -1.279 | $C_9H_{14}N_3O_2^-$ | Hercynine | [M-H] |
| 180.1027 | -1.682 | $C_{10}H_{14}NO_2^-$ | 3,4-Dimethoxyphenethylamine | [M-H] |

| | | | | |
|---|---|---|---|---|
| 208.1091 | -0.245 | $C_{10}H_{14}N_3O_2^-$ | 4-[Methyl(nitroso)amino]-1-(3-pyridinyl)-1-butanol | [M-H] |
| 224.104 | -0.294 | $C_{10}H_{14}N_3O_3^-$ | 4-(methylnitrosamino)-1-(3-pyridyl-n-oxide)-1-butanol | [M-H] |
| 183.1137 | -1.103 | $C_9H_{15}N_2O_2^-$ | N-(3-acetamidopropyl)pyrrolidin-2-one | [M-H] |
| 195.1137 | -1.035 | $C_{10}H_{15}N_2O_2^-$ | 2-Isopropyl-3,5-dimethoxy-6-methylpyrazine | [M-H] |
| 223.1085 | -1.420 | $C_{11}H_{15}N_2O_3^-$ | Butalbital | [M-H] |
| 201.1358 | 0.498 | $C_8H_{17}N_4O_2^-$ | N,N-dimethylarginine | [M-H] |
| 209.1293 | -1.205 | $C_{11}H_{17}N_2O_2^-$ | Cyclo(leucylprolyl) | [M-H] |
| 211.145 | -0.956 | $C_{11}H_{19}N_2O_2^-$ | 1,4-Bipiperidine-1-carboxylic acid | [M-H] |
| 198.1611 | -0.433 | $C_{10}H_{20}N_3O^-$ | Diethylcarbamazine | [M-H] |
| 228.1716 | -0.661 | $C_{11}H_{22}N_3O_2^-$ | N-{3-[(4-Acetamidobutyl)amino]propyl}acetamide | [M-H] |
| 311.1685 | -0.450 | $C_{17}H_{27}O_3S^-$ | 4-Undecylbenzenesulfonic acid | [M-H] |
| 299.2014 | -0.849 | $C_{20}H_{27}O_2^-$ | Tretinoin | [M-H] |
| 253.2174 | 0.380 | $C_{16}H_{29}O_2^-$ | Palmitelaidic acid | [M-H] |
| 325.1839 | -1.199 | $C_{18}H_{29}O_3S^-$ | 4-Dodecylbenzenesulfonic acid | [M-H] |
| 303.2327 | -0.837 | $C_{20}H_{31}O_2^-$ | Icosa-5,8,11,14-tetraenoic acid | [M-H] |
| 269.2488 | 0.728 | $C_{17}H_{33}O_2^-$ | Margaric acid | [M-H] |
| 305.2483 | -0.996 | $C_{20}H_{33}O_2^-$ | eicosa-8,11,14-trienoic acid | [M-H] |
| 283.2642 | -0.190 | $C_{18}H_{35}O_2^-$ | Octadecanoic acid/ Stearic acid | [M-H] |
| 766.5398 | 0.745 | $C_{43}H_{77}NO_8P^-$ | 1-stearoyl-2-arachidonoyl-sn-glycero-3-phosphoethanolamine (PE38:4) | [M-H] |
| 744.555 | 0.163 | $C_{41}H_{79}NO_8P^-$ | 1-stearoyl-2-oleoyl-sn-glycero-3-phosphoethanolamine zwitterion | [M-H] |
| 883.5344 | 0.221 | $C_{47}H_{80}O_{13}P^-$ | 1-oleoyl-2-arachidonoyl-sn-glycero-3-phospho-1D-myo-inositol | [M-H] |
| 885.549 | -0.965 | $C_{47}H_{82}O_{13}P^-$ | 1-stearoyl-2-arachidonoyl-sn-glycero-3-phospho-1D-myo-inositol (PI 38:4) | [M-H] |

**Table S3.** Processed data normality tests

| Variable | Shapiro-Wilk Statistic | Shapiro-Wilk p-value | Kolmogorov-Smirnov Statistic | Kolmogorov-Smirnov p-value |
|---|---|---|---|---|
| 127.0039 | 0.766 | 0 | 0.219 | 0.005 |
| 135.031 | 0.191 | 0 | 0.454 | 0 |
| 167.0208 | 0.67 | 0 | 0.25 | 0.001 |
| 85.0292 | 0.494 | 0 | 0.297 | 0 |
| 133.0145 | 0.879 | 0 | 0.158 | 0.086 |
| 149.0093 | 0.82 | 0 | 0.232 | 0.002 |
| 129.0192 | 0.71 | 0 | 0.24 | 0.001 |
| 172.9913 | 0.789 | 0 | 0.223 | 0.004 |
| 119.0171 | 0.874 | 0 | 0.136 | 0.193 |
| 115.0512 | 0.379 | 0 | 0.365 | 0 |
| 83.0499 | 0.393 | 0 | 0.393 | 0 |
| 115.0401 | 0.69 | 0 | 0.248 | 0.001 |
| 111.045 | 0.677 | 0 | 0.273 | 0 |
| 159.0301 | 0.859 | 0 | 0.169 | 0.053 |
| 203.0196 | 0.413 | 0 | 0.351 | 0 |
| 183.0119 | 0.379 | 0 | 0.43 | 0 |
| 130.0508 | 0.394 | 0 | 0.358 | 0 |
| 105.0378 | 0.846 | 0 | 0.18 | 0.033 |
| 177.0402 | 0.651 | 0 | 0.284 | 0 |
| 121.0658 | 0.227 | 0 | 0.479 | 0 |
| 241.0115 | 0.555 | 0 | 0.343 | 0 |
| 168.0431 | 0.786 | 0 | 0.217 | 0.005 |
| 179.0558 | 0.582 | 0 | 0.3 | 0 |
| 259.0224 | 0.538 | 0 | 0.315 | 0 |
| 299.2014 | 0.49 | 0 | 0.332 | 0 |
| 253.2174 | 0.44 | 0 | 0.371 | 0 |
| 303.2327 | 0.533 | 0 | 0.333 | 0 |
| 269.2488 | 0.558 | 0 | 0.331 | 0 |
| 283.2642 | 0.752 | 0 | 0.228 | 0.003 |

| | | | | |
|---|---|---|---|---|
| 766.5398 | 0.36 | 0 | 0.409 | 0 |
| 885.549 | 0.576 | 0 | 0.31 | 0 |
| 206.9961 | 0.458 | 0 | 0.343 | 0 |
| 130.0653 | 0.425 | 0 | 0.396 | 0 |
| 173.0269 | 0.47 | 0 | 0.335 | 0 |
| 86.0966 | 0.563 | 0 | 0.325 | 0 |
| 184.0736 | 0.694 | 0 | 0.271 | 0 |
| 255.2328 | 0.736 | 0 | 0.232 | 0.002 |
| 279.2326 | 0.457 | 0 | 0.381 | 0 |
| 419.2574 | 0.589 | 0 | 0.299 | 0 |
| 437.2675 | 0.516 | 0 | 0.392 | 0 |
| 464.3142 | 0.407 | 0 | 0.425 | 0 |
| 480.3098 | 0.525 | 0 | 0.364 | 0 |
| 616.4721 | 0.506 | 0 | 0.405 | 0 |
| 905.3844 | 0.207 | 0 | 0.476 | 0 |
| 718.5382 | 0.415 | 0 | 0.367 | 0 |
| 994.4794 | 0.686 | 0 | 0.248 | 0.001 |
| 861.5491 | 0.385 | 0 | 0.367 | 0 |
| 910.5677 | 0.801 | 0 | 0.206 | 0.009 |
| 833.5926 | 0.867 | 0 | 0.203 | 0.011 |
| 813.6008 | 0.679 | 0 | 0.292 | 0 |
| 711.6513 | 0.537 | 0 | 0.373 | 0 |
| 810.6464 | 0.805 | 0 | 0.206 | 0.01 |
| 1013.4918 | 0.776 | 0 | 0.216 | 0.006 |
| 542.4906 | 0.387 | 0 | 0.373 | 0 |

**Table S4.** Results for classification Models

| Model | Set | accuracy | f1 score | precision | recall | AUC |
|---|---|---|---|---|---|---|
| Logistic Regression | Test | 0.958 ± 0.052 | 0.953 ± 0.060 | 0.967 ± 0.067 | 0.950 ± 0.100 | 0.955 ± 0.056 |
| Logistic Regression | Train | 1.000 ± 0.000 | 1.000 ± 0.000 | 1.000 ± 0.000 | 1.000 ± 0.000 | 1.000 ± 0.000 |
| Random Forest | Test | 0.920 ± 0.098 | 0.933 ± 0.082 | 0.886 ± 0.140 | 1.000 ± 0.000 | 0.920 ± 0.098 |
| Random Forest | Train | 1.000 ± 0.000 | 1.000 ± 0.000 | 1.000 ± 0.000 | 1.000 ± 0.000 | 1.000 ± 0.000 |
| SVM | Test | 0.858 ± 0.135 | 0.862 ± 0.122 | 0.867 ± 0.163 | 0.870 ± 0.108 | 0.855 ± 0.135 |
| SVM | Train | 0.854 ± 0.035 | 0.858 ± 0.033 | 0.842 ± 0.044 | 0.875 ± 0.027 | 0.854 ± 0.034 |
| XGBoost | Test | 0.853 ± 0.083 | 0.861 ± 0.082 | 0.846 ± 0.130 | 0.920 ± 0.160 | 0.860 ± 0.080 |
| XGBoost | Train | 0.881 ± 0.061 | 0.877 ± 0.079 | 0.894 ± 0.098 | 0.895 ± 0.163 | 0.879 ± 0.064 |

**Table S5.** Logistic regression average coefficients and statistical significance. Rows which are highlighted in yellow, indicates variable (m/z) that were deemed significant ($p$ <0.05) and displayed > x2 fold change from Volcano plot. Column highlighted in grey, shows variables that displayed statistical significance ($p$ <0.05).

| Variable (*m/z*) | Assignment | Adduct | Mean Coefficient | Std Coefficient | P-Value |
|---|---|---|---|---|---|
| 127.0039 | '5-Hydroxy-2-furoic acid' | [M-H] | 1.96E-05 | 1.28E-05 | 0.026 |
| 135.031 | 'Hypoxanthine' | [M-H] | 2.06E-06 | 8.65E-06 | 0.623 |
| 167.0208 | 'Urate/ Uric Acid' | [M-H] | 0.00014 | 9.90E-05 | 0.034 |
| 85.0292 | '3-Butenoic acid/ Biacetyl/ butyrolactone/ oxolan-3-one' | [M-H] | -0.0003 | 5.37E-05 | 0 |
| 133.0145 | 'Malate/ (±)-Malic Acid' | [M-H] | -0.000215 | 6.78E-05 | 0.002 |
| 149.0093 | '(.+/-.)-Tartaric acid' | [M-H] | 8.95E-07 | 4.82E-07 | 0.014 |
| 129.0192 | 'Itaconate/ Mesaconate/ (Z)-glutaconic acid/ 4,5-dioxovaleric acid' | [M-H] | -0.00018 | 6.16E-05 | 0.002 |
| 172.9913 | Phenol sulfate' | [M-H] | -3.63E-05 | 3.23E-05 | 0.065 |
| 119.0171 | '4-Thiapentanoic acid' | [M-H] | -0.000345 | 5.12E-05 | 0 |
| 115.0512 | '1-Nitroso-3-pyrrolidinol/ Succinamide' | [M-H] | -9.27E-06 | 3.95E-06 | 0.006 |
| 83.0499 | 'Cyclopentanone' | [M-H] | -1.93E-06 | 8.71E-06 | 0.646 |
| 115.0401 | '3-Methyl-2-oxobutanoic acid/ Levulinic acid' | [M-H] | -0.000141 | 3.10E-05 | 0 |
| 111.045 | 'Sorbic acid' | [M-H] | -0.000132 | 2.98E-05 | 0 |
| 159.0301 | '2-Oxoadipate' | [M-H] | 4.34E-05 | 1.57E-05 | 0.003 |

| | | | | | |
|---|---|---|---|---|---|
| 203.0196 | '2,6-Anhydro-6-carboxy-3-deoxyhex-2-enonic acid' | [M-H] | -1.35E-05 | 1.90E-06 | 0 |
| 183.0119 | '4-styrenesulfonic acid' | [M-H] | -8.36E-06 | 1.57E-05 | 0.299 |
| 130.0508 | '5-Aminolevulinate/ propionylglycine/ Ac-Ala-OH/ Aminolevulinic acid/ L-Hydroxyproline' | [M-H] | -4.25E-05 | 8.56E-06 | 0 |
| 105.0378 | 'methionol' | [M-H] | 3.97E-05 | 1.95E-05 | 0.01 |
| 177.0402 | 'L-Gulono-14-lactone/ D-Glucono-14-lactone' | [M-H] | -0.000103 | 2.38E-05 | 0 |
| 121.0658 | 'Phenylethyl alcohol' | [M-H] | 1.14E-05 | 2.22E-05 | 0.315 |
| 241.0115 | '1D-myo-Inositol 1,2-cyclic phosphate' | [M-H] | -0.000252 | 0.000107 | 0.006 |
| 168.0431 | 'P-DMEA' | [M-H] | 0.000383 | 7.99E-05 | 0 |
| 179.0558 | 'myo-Inositol/ Fructose/ Glucose/ Galactose/ beta-D-Galactopyranose' | [M-H] | -0.00046 | 5.76E-05 | 0 |
| 259.0224 | 'D-Fructose 6-phosphate/ D-glucose 6-phosphate/ D-glucopyranose 6-phosphate' | [M-H] | -3.50E-05 | 1.16E-05 | 0.002 |
| 299.2014 | 'Tretinoin' | [M-H] | -2.64E-05 | 8.71E-06 | 0.002 |
| 253.2174 | 'Palmitelaidic acid' | [M-H] | 3.10E-05 | 1.46E-05 | 0.008 |
| 303.2327 | ' Icosatetraenoic acid | [M-H] | 3.52E-05 | 3.31E-05 | 0.076 |
| 269.2488 | 'Margaric acid' | [M-H] | 1.14E-05 | 3.14E-06 | 0.001 |
| 283.2642 | 'Octadecanoic acid/ Stearic acid' | [M-H] | -6.62E-05 | 2.35E-05 | 0.003 |
| 766.5398 | '1-stearoyl-2-arachidonoyl-sn-glycero-3-phosphoethanolamine' | [M-H] | 1.27E-05 | 8.06E-06 | 0.024 |
| 885.549 | '1-stearoyl-2-arachidonoyl-sn-glycero-3-phospho-1D-myo-inositol' | [M-H] | -0.00013 | 5.96E-05 | 0.008 |
| 206.9961 | 'Pyrogallol-2-O-sulphate' | [M+H] | -1.43E-05 | 1.14E-05 | 0.048 |
| 130.0653 | 'leucoline' | [M+H] | -4.23E-05 | 2.18E-05 | 0.012 |
| 173.0269 | 'p-Toluenesulfonic acid' | [M+H] | -0.000171 | 0.00012 | 0.032 |
| 86.0966 | Piperidine | [M+H] | -7.12E-05 | 3.23E-05 | 0.007 |
| 184.0736 | 'Choline phosphate' | [M+H] | -5.14E-05 | 8.69E-05 | 0.256 |
| 255.2328 | 'FA 16:0' | [M-H] | 0.000336 | 7.37E-05 | 0 |
| 279.2326 | 'FA 18:2' | [M-H] | -0.000229 | 8.60E-05 | 0.003 |
| 419.2574 | 'LPA O-18:2' | [M-H] | -2.70E-05 | 1.93E-05 | 0.035 |
| 437.2675 | 'LPA 18:0/LPA O-18:1;O' | [M-H] | -2.21E-06 | 3.16E-06 | 0.193 |
| 464.3142 | 'LPC O-15:1/LPE O-18:1' | [M-H] | -6.32E-06 | 3.80E-06 | 0.02 |
| 480.3098 | 'LPC 15:0/LPC O-15:1;O/LPE 18:0/LPE O-18:1;O' | [M-H] | 1.87E-05 | 4.57E-06 | 0 |
| 616.4721 | 'CerP 34:1;O2/LPC O-26:2/LPE O-29:2' | [M-H] | 3.53E-06 | 2.07E-06 | 0.019 |
| 905.3844 | 'PIP 33:7;O' | [M-H] | 0.000204 | 0.000134 | 0.027 |
| 718.5382 | 'CerP 39:1;O4/LPC 31:1;O/LPE 34:1;O/LPS O-33:1/PC 31:0/PC O-31:1;O/PE 34:0/PE O-34:1;O' | [M-H] | 2.66E-06 | 3.25E-06 | 0.141 |
| 994.4794 | 'MIPC 34:6;O6' | [M-H] | 2.96E-05 | 1.13E-05 | 0.004 |
| 861.5491 | 'PI 36:2/PI O-36:3;O' | [M-H] | -6.83E-05 | 3.92E-05 | 0.017 |
| 910.5677 | 'PI-Cer 39:2;O6' | [M-H] | 2.25E-05 | 7.18E-06 | 0.002 |
| 833.5926 | 'PG 39:1;O' | [M-H] | -0.000208 | 0.000134 | 0.025 |
| 813.6008 | 'PA 43:2;O/PG O-40:3' | [M-H] | 1.13E-05 | 8.77E-06 | 0.045 |
| 711.6513 | 'DG O-41:0;O2' | [M-H] | 7.64E-06 | 6.58E-06 | 0.06 |
| 810.6464 | 'HexCer 41:2;O3' | [M-H] | 1.13E-05 | 1.11E-05 | 0.085 |
| 1013.492 | 'PIP 39:5' | [M+Cl] | 3.63E-05 | 2.08E-05 | 0.017 |

| | | | | | |
|---|---|---|---|---|---|
| 542.4906 | 'Cer 34:2;O/NAE 32:2' | [M+Na] | 3.66E-05 | 2.94E-05 | 0.049 |

**Table S6.** Molecules present on the surface of the deposit (close to the interface) categorised as either interfacial or distant from interface at 1 day and 28 days

| **1 day** | **28 days** |
|---|---|
| **Interfacial** | **Interfacial** |
| • Succinic anhydride<br>• Xylonic acid<br>• myo-Inositol/ Fructose/ Glucose/ Galactose/ Galactopyranose<br>• Itaconate/ Mesaconate/ Glutaconic acid/ 4,5-dioxovaleric acid<br>• Inositol cyclic phosphate<br>• *PIP 39:5*<br>• *FA 18:2*<br>• *PI 38:4*<br>• *FA 18:1*<br>• *FA 18:0*<br>• *LPA O-18:2*<br>• *Eicosatetraenoic acid*<br>• *FA 16:0*<br>• *Hex2Cer 34:4;O4* | • Inositol cyclic phosphate<br>• Cytosine<br>• Thymine<br>• Uracil<br>• Adenine<br>• Guanine<br>• L-Proline<br>• *FA 16:0*<br>• *FA 18:1*<br>• *FA 18:0*<br>• *Eicosatetraenoic acid*<br>• *LPA O-18:2*<br>• *Hex2Cer 34:4;O4*<br>• *PI 38:4* |
| **Distant from interface** | **Distant from interface** |
| • Cytosine<br>• Thymine<br>• Uracil<br>• Adenine<br>• Guanine<br>• L-Proline<br>• *SHexCer 41:5;O5*<br>• *PG 39:1;O* | • *SHexCer 41:5;O5*<br>• *PG 39:1;O*<br>• *PIP 39:5*<br>• *FA 18:2* |

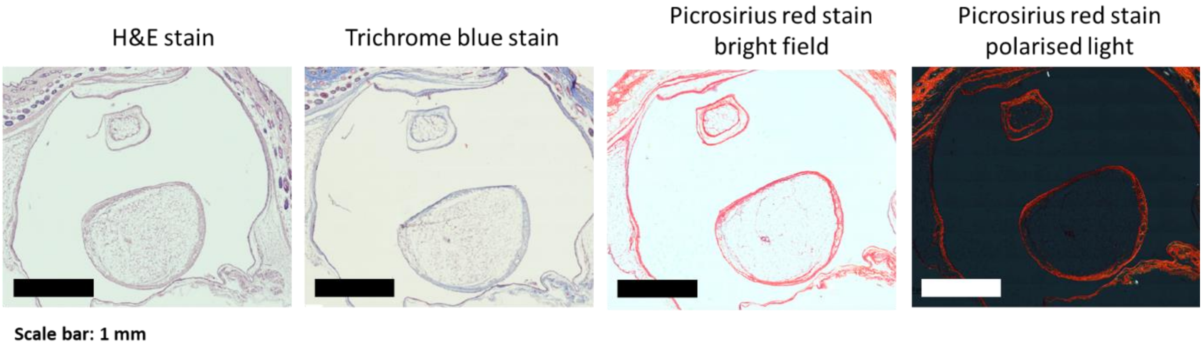


**Figure S1.** The histological stain of the surrounding tissue. The tissue were stained with Haematoxylin and eosin (H&E), Masson's trichrome stain and picrosirius red stain

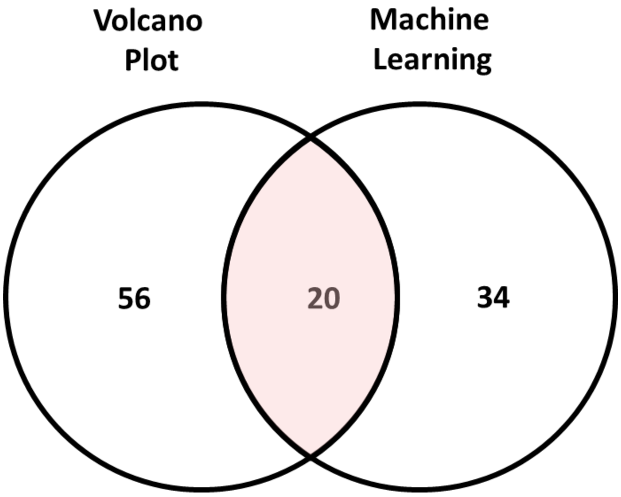


**Figure S2.** The Venn diagram shows the number of assignments overlap between the logistic regression analysis with the univariate statistics Volcano plot approach.

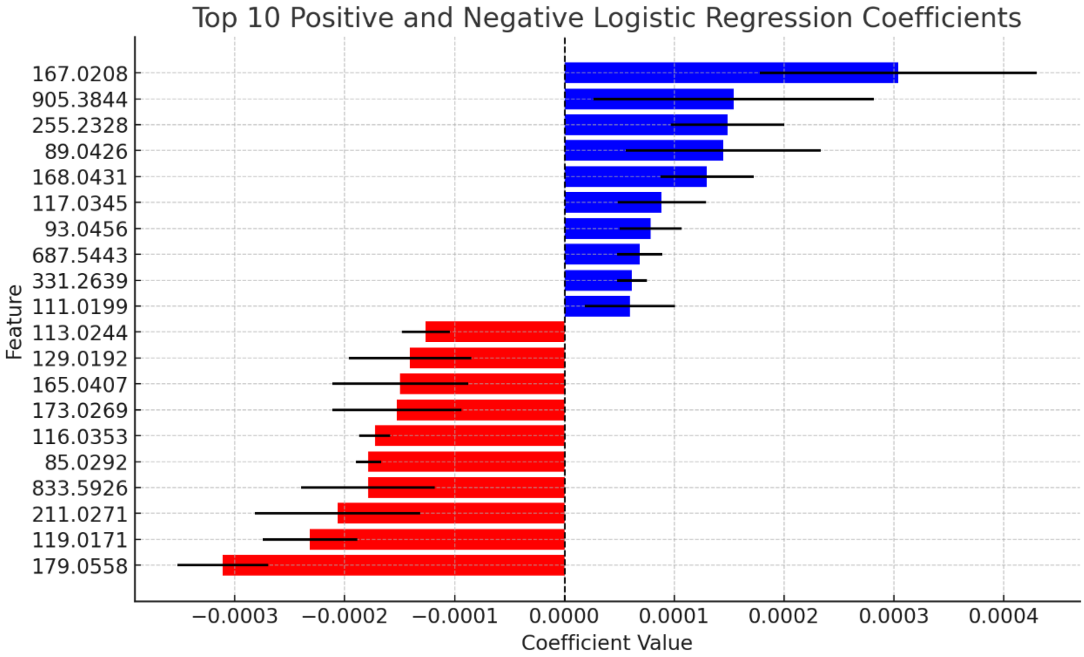


**Figure S3.** Top ten positive and top ten negative logistic regression coefficients

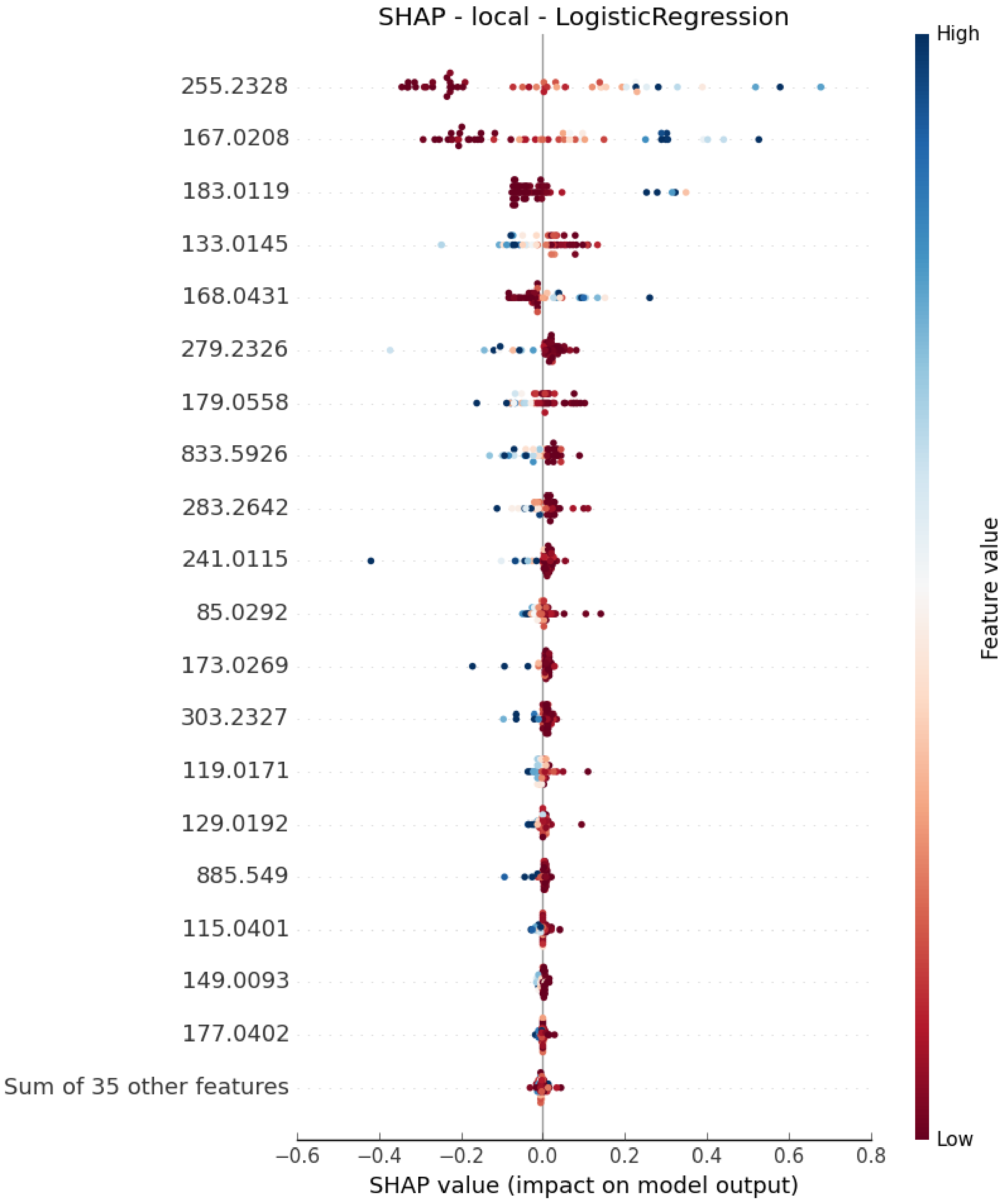


**Figure S4.** Shapley (SHAP) importance values for Logistic Regression

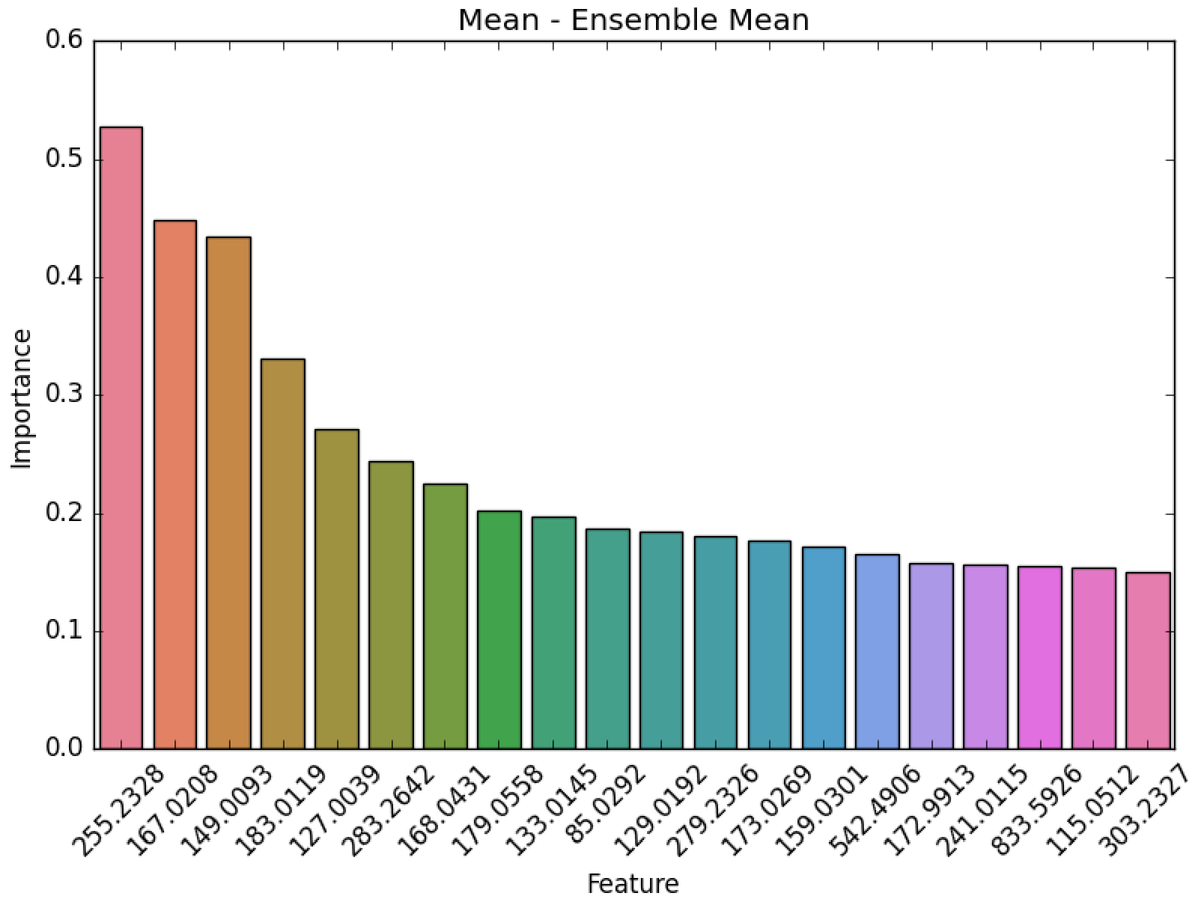


**Figure S5.** Average feature importance across modelling approaches

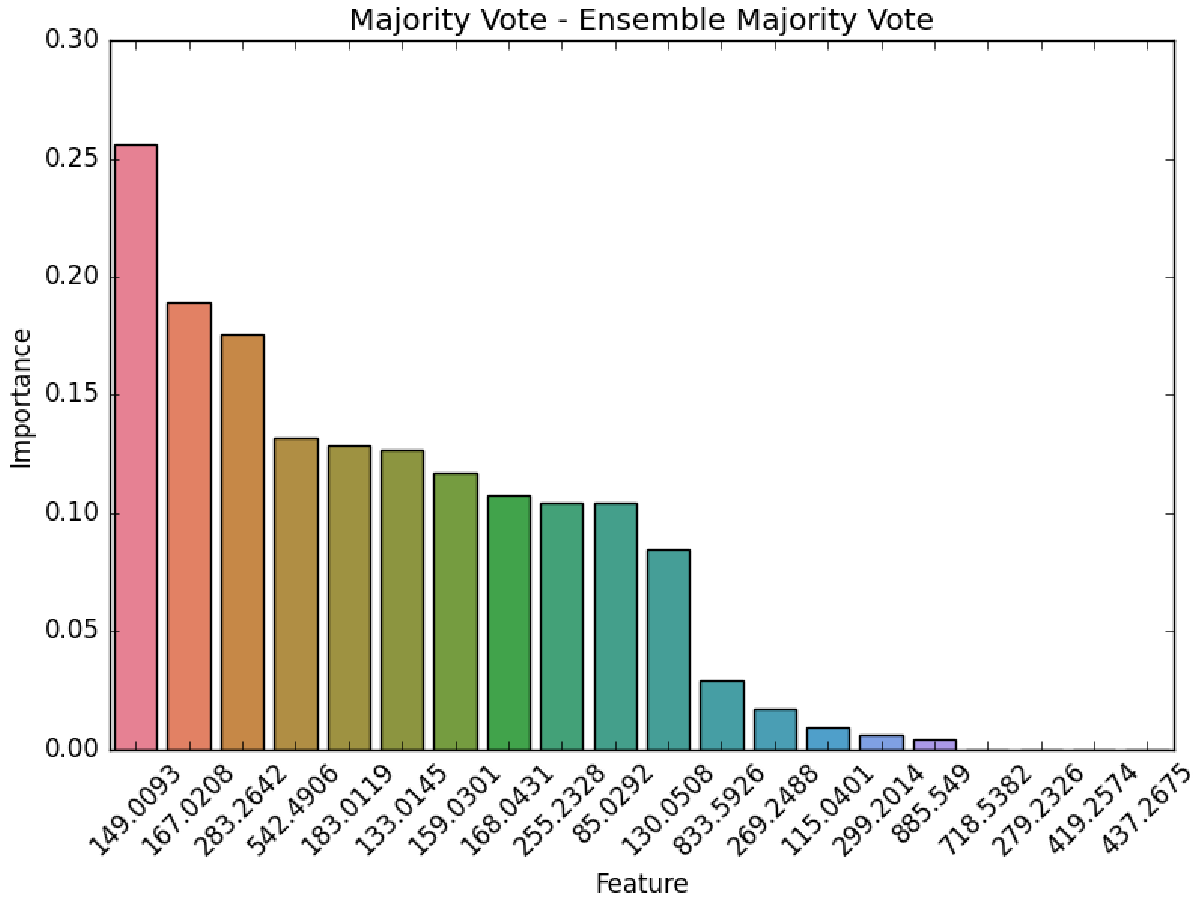


**Figure S6.** Majority voting results determining feature importance across modelling approaches

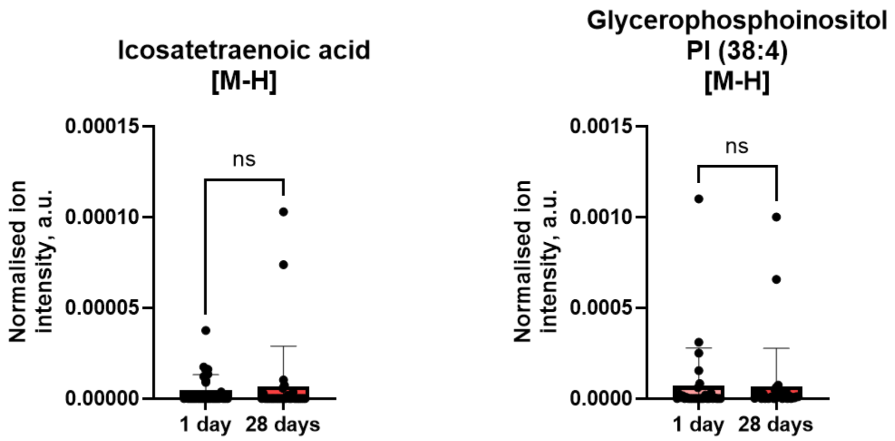


**Figure S7.** Bar charts of normalised ion intensity of icosatetraenoic acid and glycerophosphoinositol PI (38:4). Data are plotted as normalised ion intensities to total ion count. Data are displayed as n=30, mean ± SD.

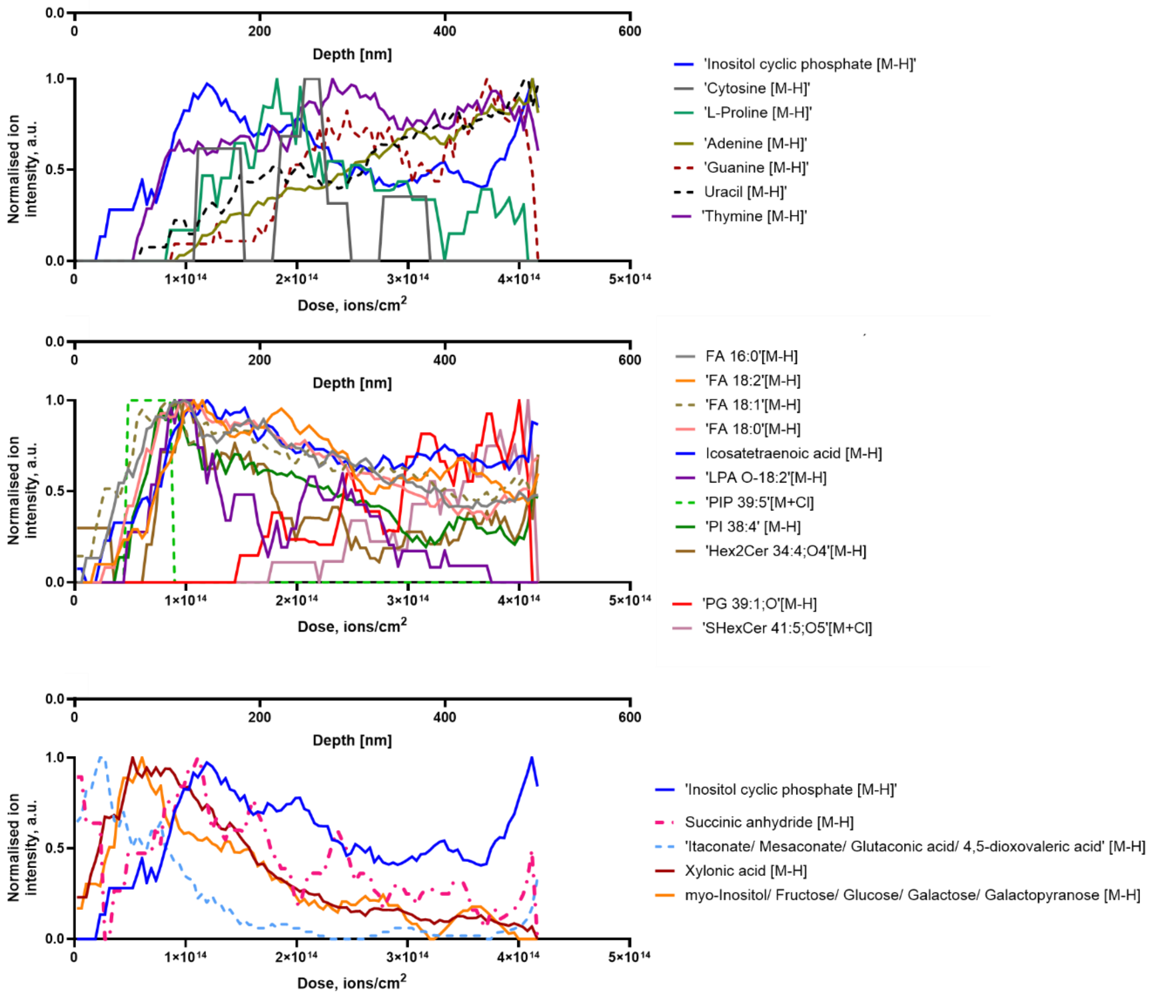


**Figure S8.** Depth profile at silicone catheter surfaces versus primary ions dose and estimated depth displaying lipids, amino acid, nucleic acid bases and sugar-based metabolites molecular ions profile at 1 day.

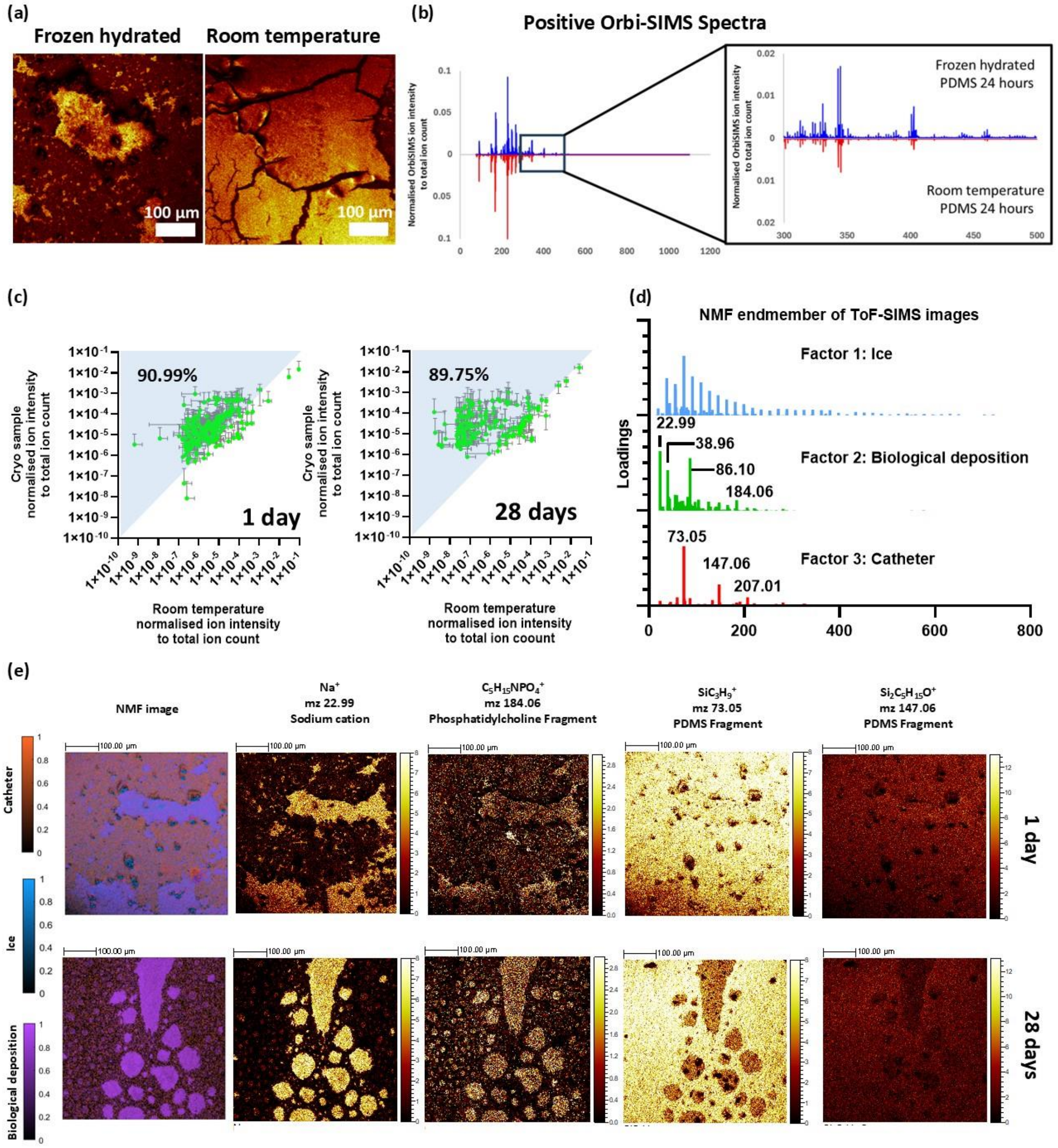


**Figure S9** (a) LMIG/ToF-SIMS ion images of the catheter surfaces when sample were analysed under frozen hydrated and room temperature conditions. (b) Comparison of GCIB/OrbiSIMS spectra of catheter surfaces analysed under frozen-hydrated (blue) and room temperature (red) conditions. (c) Scatter plot comparing the normalised ion intensities of biomolecules detected when the surface of the catheters was analysed under room temperature and frozen hydrated. (d) Non-negative matrix factor (NMF) loadings showing the separation of LMIG/ToF-SIMS data into three distinct components: ice, biological deposition, and catheter with (e) representative NMF images for 1- and 28-days implantation following 500 NMF iterations. The NMF analysis was performed using loadings and images from raw LMIG/ToF datasets. Colour scale represents the intensity of NMF endmembers post calculation along with the ion images for C5H15NPO4+ which represent phosphatidylcholine fragments and SiC3H9+ and Si2C5H15O+ which is used as a marker for PDMS. Scale bar: 100 µm.

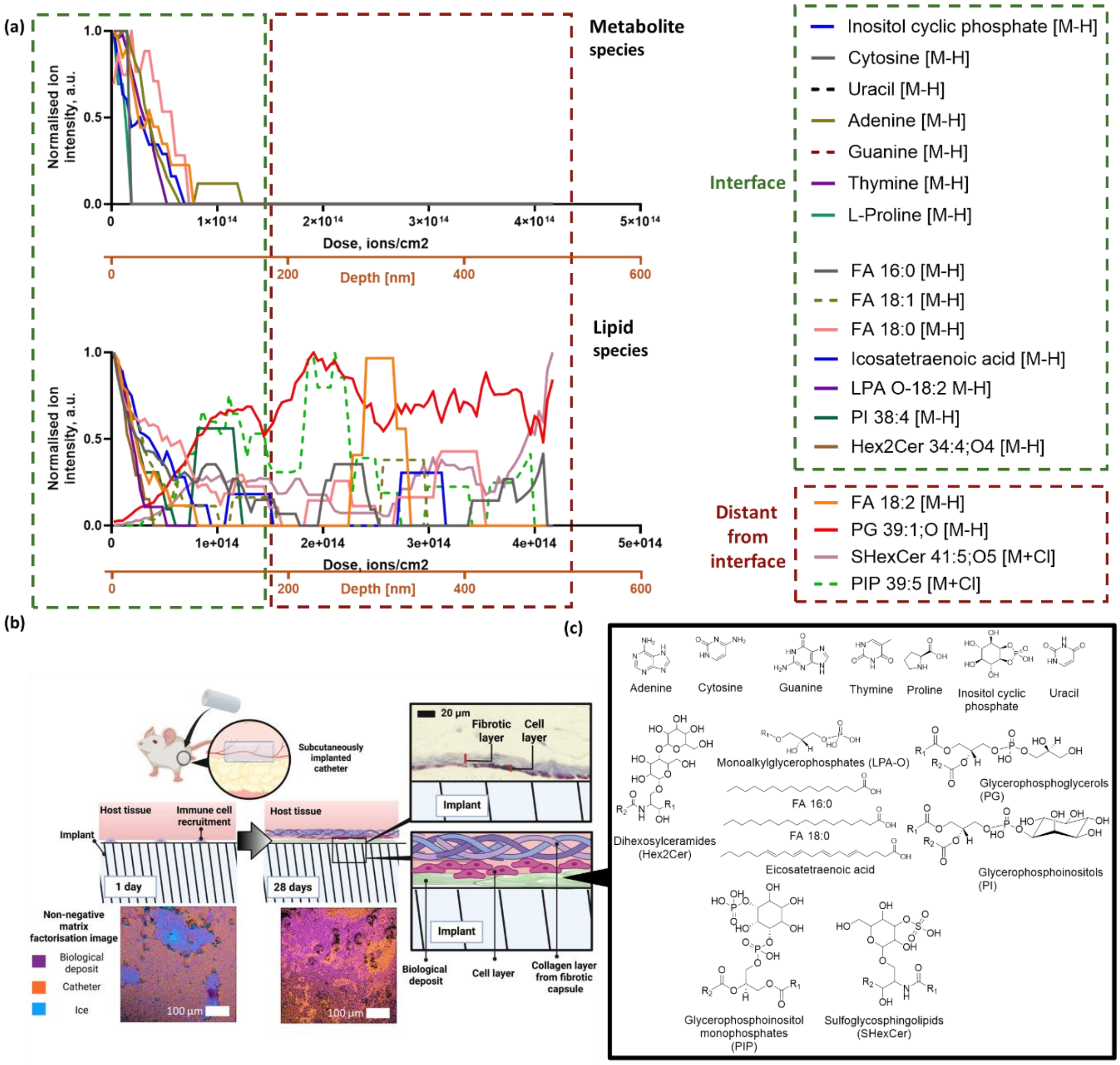


**Figure S10** Depth profile at silicone catheter surfaces versus primary ions dose (ions/cm$^2$) and estimated depth displaying lipids, amino acid and nucleic acid bases molecular ions at 28 days. Molecules present on the surface of the deposit (close to the interface) is categorised as interfacial while those present deep in the deposit is categorised as distant from interface. The secondary x axis presents depth estimated from comparison with organic standards **(b)** Schematic illustrating and summarising the presence of biological deposit formed on the surface of medical device following implantation **(c)** Chemical structure of species detected by cryo-OrbiSIMS.

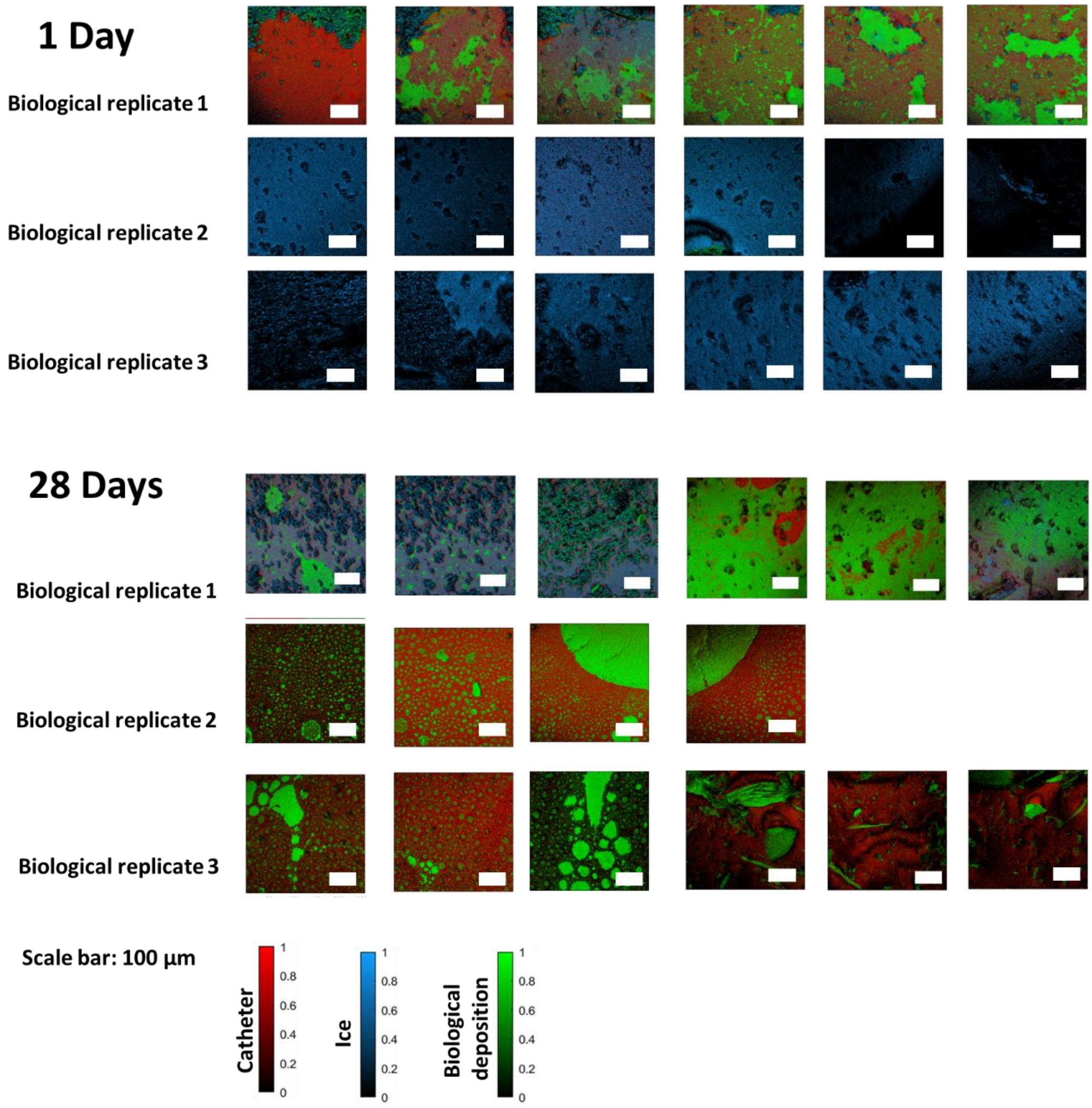


**Figure S11.** NMF Images of catheter explant collected Balb/c mice at 1 day and 28 days. Colour scale represents the intensity of NMF endmembers post calculation The absence or minimal signal from catheter or biological deposits for these NMF images for biological replicate 2 and 3 was attributed to the formation of ice on the surface of the catheter during sample analysis. The formation of ice on the surface prevents from any further ToF-SIMS data analysis, however this does not impact the Orbi-SIMS data acquisition

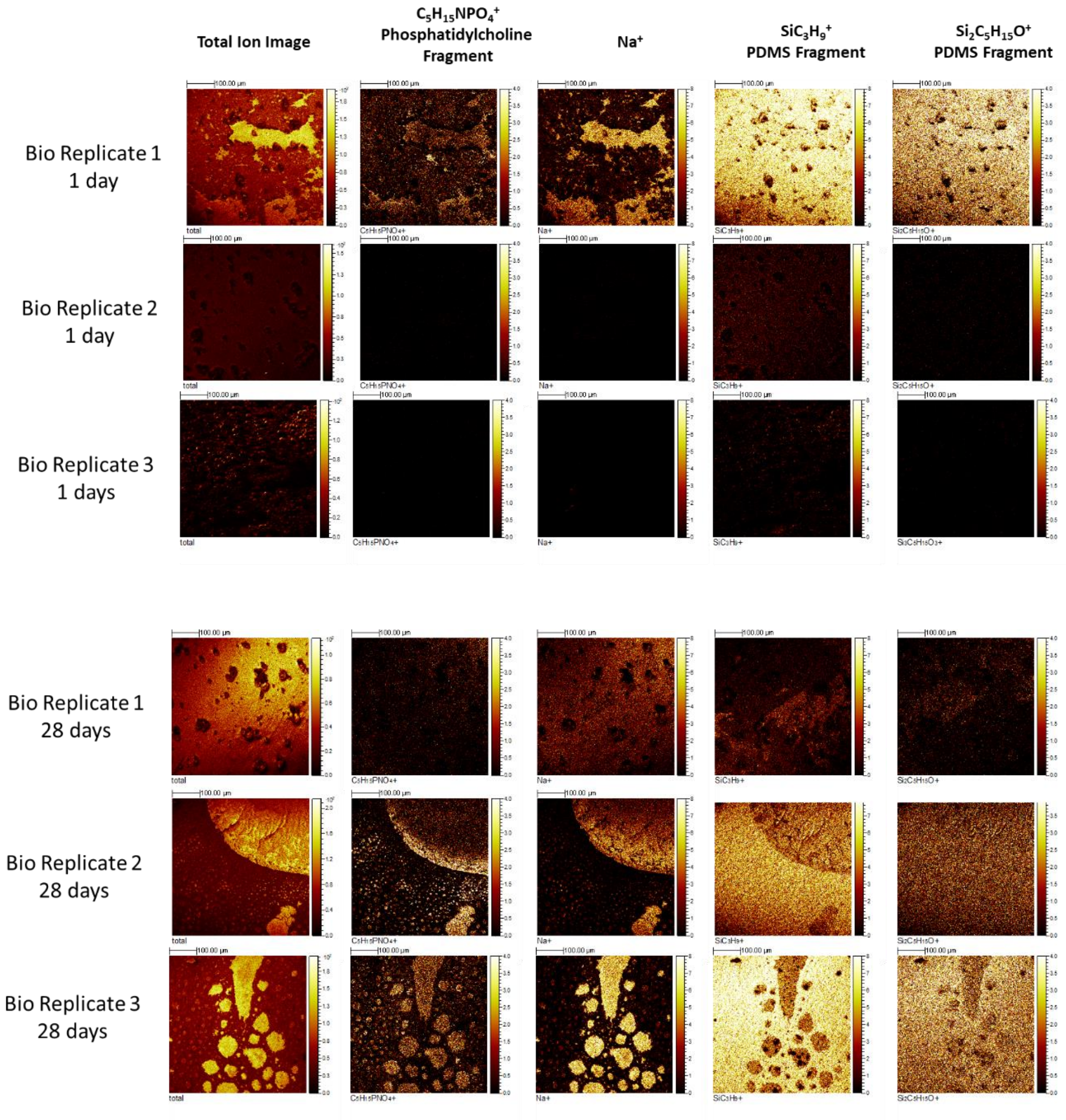


**Figure S12.** ToF-SIMS chemical ion images of catheter explant collected Balb/c mice at 1 day and 28 days. The ion images selected are $C_5H_{15}NPO_4^+$ Phosphatidylcholine fragments and $Na^+$ which are used as markers for biological deposits for ToF images and $Si_2C_5H_{15}O^+$ which is used as a marker for PDMS oligomer fragment. The absence or minimal signal for these ion images for biological replicate 2 and 3 was attributed to the formation of ice on the surface of the catheter during sample analysis. The formation of ice on the surface prevent from any further ToF-SIMS data analysis, however this does not impact the Orbi-SIMS data acquisition.
